# Laboratory transmission of Adult Salmon Enteritis and associated pathogens in juvenile Chinook Salmon (*Oncorhynchus tshawytscha*)

**DOI:** 10.1101/2025.04.07.647091

**Authors:** Tamsen Polley, Claire E. Couch, Connor Leong, James T. Peterson, Louis M. Weiss, Peter M. Takvorian, Michael L. Kent

## Abstract

Adult Salmon Enteritis (ASE), characterized by severe ulcerative enteritis, has been linked to pre-spawn mortality (PSM) in spring Chinook Salmon (*Oncorhynchus tshawytscha*) in certain rivers in Oregon, USA. Catastrophic losses of spring Chinook Salmon have resulted from PSM, a significant threat to their population stability. Understanding the causes of ASE is therefore critical for mitigating PSM and supporting conservation. This study investigates the potential infectious etiology of ASE using a juvenile Chinook Salmon model. Fish were immunocompromised with dexamethasone implants, fasted, and exposed to intestinal tissues from ASE-affected adult Chinook. Histopathology of recipient fish revealed mid-intestinal lesions consistent with ASE. The microsporidium *Enterocytozoon schreckii*, which is observed in ASE-affected adults from rivers, was transmitted for the first time to juvenile Chinook Salmon, making *E. schreckii* a potential new pathogen of juvenile salmon. Additionally, intranuclear inclusions were identified in enterocytes by histopathology and viral particles were detected by electron microscopy in recipient fish. The study demonstrates that intestinal lesions consistent with ASE can be experimentally induced in juvenile Chinook Salmon through oral exposure to infected tissues, supporting an infectious etiology. Further research is needed to isolate specific pathogens, including viruses and *E. schreckii*, and to elucidate their roles in ASE development.

## 1 Introduction

Chinook Salmon (*Oncorhynchus tshawytscha*) are integral to the economic, cultural, and environmental success of the Pacific Northwest of North America and elsewhere in their natural distribution. However, Chinook salmon populations in Oregon and across their natural range are in decline due to multiple stressors, including climate change, drought, habitat loss, poor ocean conditions, overfishing, disease, and dams that block migration.

Spring Chinook Salmon are anadromous and semelparous, returning to their natal rivers in late winter through spring, completing sexual maturation during migration to spawning locations, and spawning in later summer and fall. This has allowed them to be uniquely sensitive to diverse environmental impacts after they return to freshwater to spawn due to their longer holding period. After reaching freshwater they are anorexic and become immune compromised - i.e., have extremely elevated cortisol levels and reduced innate immunity (Dolan et al., 2016; Schreck, 2000). These conditions, compounded by additional environmental stressors, place spawning salmon at high risk for disease and mortality upon returning to freshwater.

Survival during this sensitive period prior to spawning is critical for future spring Chinook Salmon populations, as these semelparous fish die after spawning. However, pre-spawning mortality (PSM)—where salmon die before spawning—has been documented at catastrophic levels in the Willamette Basin, with over 90% loss in certain runs during a single season, and its severity appears to be increasing (Carey et al., 2024; Naughton et al., 2023). PSM occurs after salmon return to their natal rivers, but prior to spawning either at the spawning grounds or en route during migration. While the underlying cause of PSM in spring Chinook Salmon remains elusive, various factors have been associated with this phenomenon. These include anthropogenic stressors such as transport over dams (Keefer et al., 2010); environmental conditions including stream temperature and stocking density (Bowerman et al., 2021; Keefer et al., 2010; Roumasset, 2012); individual factors including wild versus hatchery origin and fish length (Bowerman et al., 2021); and pathogen burdens or lesions (Benda et al., 2015; Nervino et al., 2024).

Since 2009, we have used necropsy and histopathology to investigate PSM in spring Chinook Salmon in the Willamette Basin (hereafter, Chinook Salmon) (Benda et al., 2015; Colvin et al., 2015; Kent et al., 2013; Nervino et al., 2024). During these investigations, we identified a previously unrecognized intestinal disease that is strongly correlated with PSM in affected fish (Couch et al., 2022; Nervino et al., 2024). This disease, which we define as Adult Salmon Enteritis (ASE), is characterized by severe ulcerative enteritis, including extensive epithelial loss and diffuse inflammation of the intestinal lamina propria, in Chinook salmon from the Willamette River watershed in Oregon, USA. To date, ASE has only been observed in adult Chinook Salmon after returning to freshwater to spawn.

Initially, ASE in PSM fish was thought to result from accelerated senescence, a process expected in semelparous species (Nervino et al., 2024). However, our recent survey of Chinook Salmon populations in Oregon and Washington revealed that ASE was entirely absent in two Washington spawning populations but present in four Oregon rivers. This geographic disparity, combined with the severity of the lesions, suggests an alternative hypothesis: ASE may have an infectious etiology. Previous investigations evaluated two intestinal pathogens, *Ceratonova shasta* (Myxozoa) and *Enterocytozoon schreckii* (Microsporidia), but neither correlated with ASE severity (Nervino et al., 2024). However, during the characterization of *E. schreckii* by electron microscopy (Couch et al., 2022), we observed structures consistent with viral particles within the intestine of a fish with ASE. To date, *E. schreckii* and these viral particles have only been observed in Chinook Salmon populations affected by ASE, raising the possibility that an unidentified pathogen, potentially viral, contributes to ASE development.

To test the hypothesis of an infectious etiology for ASE, we conducted *in vivo* laboratory experiments with juvenile Chinook Salmon. First, we developed a surrogate model for spawning adult Chinook Salmon using juvenile Chinook Salmon that were fasted and immune compromised with dexamethasone. A second experiment evaluated the transmission of ASE by exposing these fish to ASE-affected intestinal tissue.

## 2 Materials and Methods

### 2.1 Fish

All animal husbandry methods and experimental procedures were performed in accordance with relevant guidelines and regulations and were approved by Oregon State University (OSU) Institutional Animal Care and Use Committee (ACUP #2023-0395). We obtained 750 juvenile Chinook Salmon weighing approximately 1 g in May 2023 from Willamette Hatchery in Oakridge, Oregon, USA (43°44’50.4”N, 122°26’41.43”W). Fish were pre-screened for furunculosis (*Aeromonas salmonicida*) and bacterial gill disease (*Flavobacterium* spp.) before acquisition by Oregon State University. Fish were held at the Fish Performance and Genetics Laboratory (FPGL), OSU in Corvallis, Oregon (44°57′62.5′′N, 123°24′11.6′′W). Fish were held at a density of approximately 2 kg fish/cubic meter in 1.5 meter in diameter circular tanks before being transferred to the Aquatic Animal Health Laboratory (AAHL) for Phase 1 and Phase 2 experiments as described below. The water supply was constant flow from gravel filtered well water. Water temperature at FPGL was a constant at 12-13 °C. Fish were fed Bio-Oregon® (Longview, Washington, USA) BioVita Fry 1.2 mm pellet diet at a maintenance ration of 1.5% fish biomass every day.

### 2.2 Histopathology

A complete necropsy was conducted on selected fish that were euthanized with 200 mg/L tricaine methane sulfonate (MS-222) (Argent Laboratories, Redmond, WA, USA). Necropsies consisted of close external and internal visual examination of each fish for signs of pathologic changes. Fresh tissues, or whole fish with the coelomic cavity incised, were collected from all study groups for histology and were placed in 10% Neutral-Buffered Formalin (NBF). After 48-hour fixation, tissues (e.g., gill, gastrointestinal tract, kidney, liver, spleen, heart, swim bladder) were trimmed, placed in tissue cassettes, and submerged in 10% NBF. Tissues were processed into paraffin blocks for histopathological slides at the Oregon Veterinary Diagnostic Laboratory (OSU, Corvallis, Oregon, USA). Slides were cut at 5 μm thickness and stained with hematoxylin and eosin (HCE). We previously showed that detection of microsporidian spores is enhanced with the Luna and Giemsa stains (Peterson et al., 2011), and hence these stains were applied to intestinal sections of selected recipient fish from Phase 2.

#### Intestinal scores

Descriptions of the histopathological changes of the fish are provided, with special attention placed on the intestines. Slides were scored for selected intestinal endpoints that varied amongst the different groups based on previous observations and lesion scoring criteria (Nervino et al., 2024). Slides were assessed by only one of the authors (T.P.) to avoid inter-reader variability. Intestinal scores established in our earlier paper (Nervino et al., 2024) assessed overall intestinal mucosal architecture in fish with ASE; i.e., percent of remaining intact epithelium, percent dysplastic epithelium, and inflammation severity involving expansion of inflammatory cells in the lamina propria and extension of the inflammatory population into the epithelium, muscularis, or serosa. The percent of remaining intact epithelium in the present study, also described as ulceration, were also scored as follows: 0, 100% remaining epithelium; 1, 99-66% remaining epithelium; 2, 65-33% remaining epithelium; and 3, 32-0% remaining epithelium. Other changes in the present study were scored using a 0 to 3 system. Epithelial dysplasia was scored as follows: 0, no apparent dysplasia; 1, 1-33% epithelial dysplasia; 2, 34-66% epithelial dysplasia; and 3, 67-100% epithelial dysplasia. Inflammation of the intestine, or enteritis, was scored as follows: 0, no apparent increase in inflammatory infiltrate, where predominantly sentinel mononuclear cells were infrequent, evenly distributed, and limited to the lamina propria, forming aggregates of ten or less leukocytes, that did not expand the intestinal fold base and the stratum compactum; 1, mild increase in inflammatory infiltrate in the lamina propria with rare extension into the epithelium or muscularis, and are expanding the distance between the intestinal fold base and the stratum compactum in <33% of the folds; 2, moderate inflammation in the lamina propria with common extension into the epithelium or muscularis, and expansion between the intestinal fold base and the stratum compactum in <66% of the folds; 3, severe increase in inflammation in the lamina propria with common extension into the epithelium or muscularis, and expansion between the intestinal fold base and the stratum compactum in >66% of the folds. Intestinal fold height was scored as follows: 0, no apparent decrease in average fold height and folds extend to intersect at roughly the midpoint of the lumen; 1, <33% decrease in fold height; 2, 34-66% decrease in fold height; 3, >66% decrease in fold height.

#### Parasite scores

The presence of pathogens of interest (*E. schreckii* and *C. shasta*) were identified as described in Nervino et al. (2024) and scored as: 0, absent/not apparent; or 1, present.

#### Renal scores

Melanin deposition within interstitial melanomacrophages (renal melanosis) was scored as this has been directly correlated negative energy balance. Renal melanosis was scored as follows: 0, 0-5% of the interstitial cell population contains melanin-laden macrophages; 1, 5-20% of the interstitial cell population contains melanin-laden macrophages; 2, 21-40% of the interstitial cell population contains melanin-laden macrophages; 3, 41-60% of the interstitial cell population contains melanin-laden macrophages.

#### Liver scores

Increased intracytoplasmic vacuolar changes in hepatocytes are expected in fish mobilizing adipose stores or ingested lipid. Scored hepatocellular vacuolar changes included cytoplasmic microvesicular, macrovesicular, and feathery/ground glass in appearance, where lesions may be patchy to diffuse. Ordinal scores for these endpoints ranged from 0 to 3, where: 0, absent/not apparent; 1, 1-33% of the average hepatocellular cytoplasm is displaced by vacuoles; 2, 34-66% of the average hepatocellular cytoplasm is displaced by vacuoles; 3, 67-100% of the average hepatocellular cytoplasm is displaced by vacuoles.

#### Myositis and coelomitis scores

Both focal to multifocal myositis and regional coelomitis (comparable to peritonitis in mammals) were scored as: 0, absent/not apparent; or 1, present.

Histology microphotographs were collected using Zeiss software and a Leica DMLB microscope using an Axiocam 105 color camera.

### 2.3 Phase 1

The purpose of Phase 1 was to determine the appropriate dexamethasone (Dex) and fasting regimen to use in our ASE exposure study (Phase 2). After approximately 28 days of acclimation at the FPGL, the fish were transported to the adjacent AAHL, where they were held in outdoor tanks with UV-sterilized water (2 L·min^-1^·tank^-1^) from the same groundwater source as the FPGL water. Water temperature at the AAHL fluctuated between 10.8–13.4°C due to seasonal temperature changes in the water supply. During the acclimation period, fish were fed the same diet as at the FPGL.

Fish were divided into four 380-liter circular tanks with 30 fish/tank or two 700-liter circular tanks with 80 to 85 fish·tank^-1^, respectively. They were allowed to acclimate for an additional 28 days, and then placed on experimental diets and administered with Dex implants (Table 1). Fish were anesthetized prior to administering Dex implants and all subsequent handling events by immersion in 50 mg/L tricaine methane sulfonate (MS-222) buffered to neutrality with sodium bicarbonate.

**Table 1.**
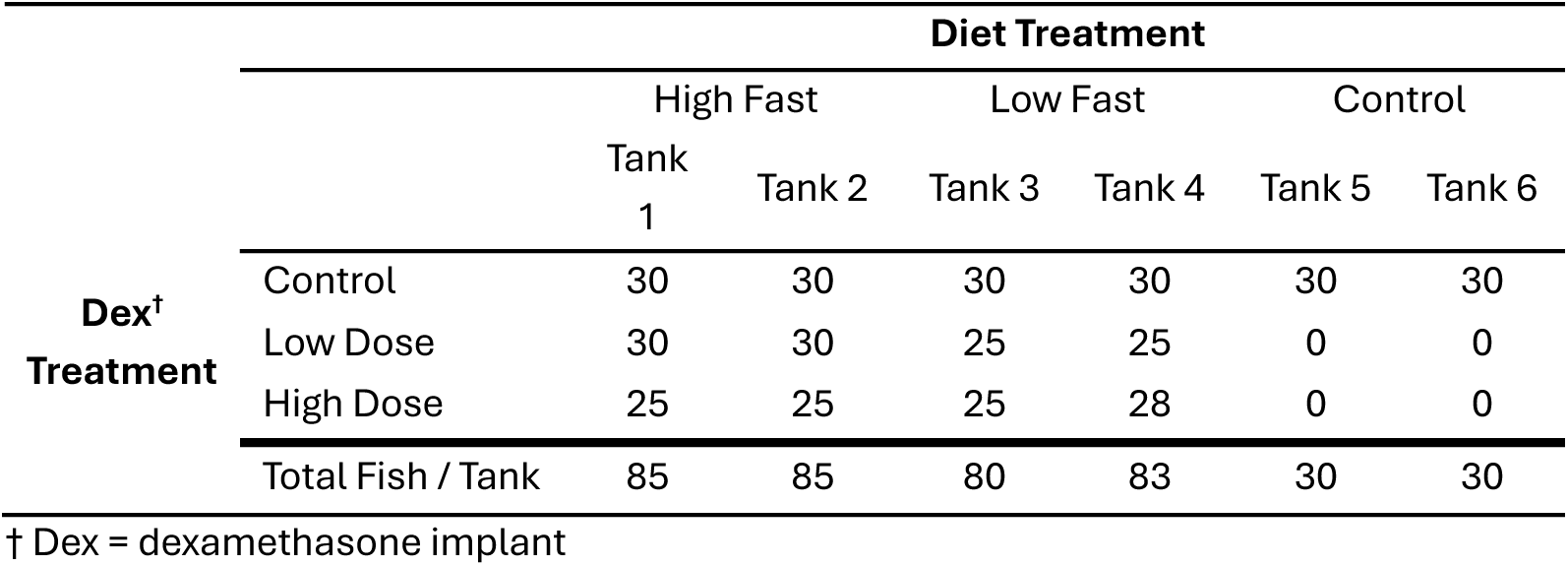
**Phase 1: Experimental design** for fasting and immunosuppression dose. A total of 393 juvenile Chinook Salmon were distributed between six 100-liter tanks in groups of 25-30. Treatments were applied in duplicates including fasting (high fast and low fast) and Dex^†^ dose (high dose and low dose).

The Dex working solution was prepared by combining 70% vegetable shortening (Crisco®) and 30% peanut oil as a vehicle with the appropriate amount of Dex powder (D4902, lot SLCP0325; Sigma-Aldrich, St. Louis, MO, USA). To prepare the solution, vegetable shortening and peanut oil were heated together until they formed a homogenous mixture. The solution was then cooled to 50°C, and the Dex power was added. After preparation, the solution was stored at room temperature in sealed containers.

Slow-release Dex implants were prepared by dissolving Dex in the vehicle so that fish would receive either 0.25 mg/g body weight (low dose) or 0.5 mg/g body weight (high dose) as a 100 μL intraperitoneal (IP) inoculum. The Dex treatment baselines were validated in our previous study with juvenile Chinook Salmon (Couch et al., 2023). The warmed (approximately 40°C) liquid-phase implants were administered by IP injection into the caudoventral abdominal cavity using 16-gauge needles. We administered Dex or control (vegetable shortening/peanut oil only) implants to each fish, which were separated into individual tanks by diet treatment group only and randomly assigned by Dex treatment group (Table 1). Within each Dex treatment group, fish were randomly assigned and marked for visual identification with a single fin clip: adipose fin for high-dose Dex, left pelvic fin for low-dose Dex, right pelvic fin for control, or no fin clip for control. Control fish (Tanks 5 and 6) were fed a normal ration and received no Dex treatment. These fish were divided into two subgroups, right pelvic fin clip with vehicle sham implant or no fin clip with no implants. These tanks were included to evaluate the effects handling and injection of the sham vehicle on survival.

Fish were recovered from anesthesia in a container with an air stone, then fish were returned to experimental tanks once appropriate independent buoyancy and swimming was observed. Fish were equally distributed in replicate 100 L tanks for each experimental diet: Bio-Oregon® BioVita Fry 1.2mm pellet diet at 3.5% fish biomass (control diet) and 1.0% fish biomass (low fasting diet) fed on Mondays, Wednesdays, and Fridays, and food withheld (high fasting diet) (Table 1).

At 24 days post-Dex treatment, all fish were treated with a 1-hour formalin bath once a day for two days at 125 ppm and 167 ppm, respectively, to control infections by the obligate parasite *Ichthyobodo necator* and subsequent colonization by the saprophyte *Saprolegnia* sp. These underlying infections were associated with increased mortality in the Dex treated fish. Both pathogens were diagnosed by experienced facility staff, where *I. necator* was confirmed by visualizing small motile flagellates on gills in wet mounts and macroscopic examinations for *Saprolegnia* sp. Additionally, 15 fish from each tank and treatment group were randomly selected for euthanasia, which was performed by submersion in 250 mg/L MS-222 in accordance with the AVMA Guidelines for the Euthanasia of Animals (Leary et al., 2020). At 50 days post-Dex treatment all remaining fish were euthanized and processed for histology as previously described. Throughout the study period, any fish that became moribund (generally ataxic accompanied by loss of buoyancy control and/or increased respiration rate) were humanely euthanized and these moribund fish were classified as mortalities.

### 2.4 Phase 2

We proceeded with Phase 2 (ASE transmission) using results of Phase 1 as guidelines – i.e., data from Section 3.1.1 was used to determine diet and Dex treatment parameters for baseline immunosuppressed and fasted fish for Phase 2. After approximately 12 weeks, 240 of the fish being held at the AAHL were transported to indoor tanks with the same parameters as 2.1.1 Phase 1. Fish were divided into eight 100 L rectangular tanks with round edges in groups of 30 in duplicate: DexFast Only, DexFast+ASE, ASE Only, Control (Table 2). Fish were administered preventative formalin baths as described in Section 2.1.1 Phase 1 to control for *I. necator* and *Saprolegnia* sp. infections and allowed to acclimate for an additional seven days. Then fish were placed on experimental diets and administered Dex implants for the pre-exposure period of 23 days. Fish were anesthetized and recovered as described in Section 2.3 Phase 1. See Figure 1 for a detailed timeline of Phase 2.

**Table 2.** **Phase 2: Experimental design** with four groups of juvenile Chinook Salmon in duplicate distributed between eight tanks: 1) DexFast^†^ + ASE^‡^, 2) DexFast^†^ Only (DexFast^†^ without ASE^‡^), 3) ASE^‡^ Only, and 4) Control (neither DexFast^†^ nor ASE^‡^).

| Treatment Groups | Number of Fish / Tank |  |  |  |  |  |  |  | Total Fish / Group |
| --- | --- | --- | --- | --- | --- | --- | --- | --- | --- |
|  | Tank 1 | Tank 2 | Tank 3 | Tank 4 | Tank 5 | Tank 6 | Tank 7 | Tank 8 |  |
| Group 1: DexFast<sup>†</sup> + ASE<sup>‡</sup> | 0 | 0 | 30 | 0 | 0 | 0 | 30 | 0 | 60 |
| Group 2: DexFast<sup>†</sup> Only | 0 | 30 | 0 | 0 | 0 | 30 | 0 | 0 | 60 |
| Group 3: ASE<sup>‡</sup> Only | 30 | 0 | 0 | 0 | 30 | 0 | 0 | 0 | 60 |
| Group 4: Control (no treatment) | 0 | 0 | 0 | 30 | 0 | 0 | 0 | 30 | 60 |
| Total Fish / Tank | 30 | 30 | 30 | 30 | 30 | 30 | 30 | 30 | 240 |

<sup>†</sup> DexFast = fish received dexamethasone implants and were placed on fasting diet <sup>‡</sup> ASE = Adult Salmon Enteritis, exposed to gastrointestinal tissue homogenate from adult fish with ASE via oral gavage

**Figure 1.**
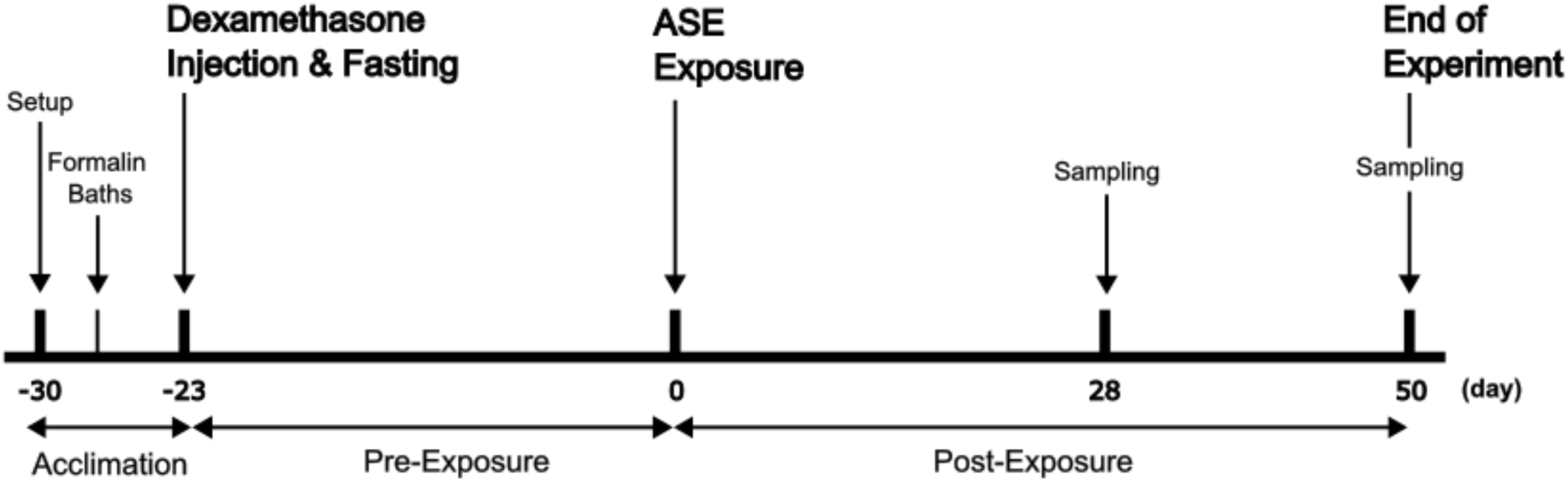
**Phase 2: Timeline of the experiment** exposing fasted and immunosuppressed juvenile Chinook Salmon to ASE-affected tissue by oral gavage. All fish were acclimated to experimental tanks for 12 weeks prior to the experiment, then were sorted into eight tanks in groups of 30 at time of experimental setup (“Setup”). On day -29 fish were treated with a formalin bath to control for external bacterial, parasitic, and water mold pathogens.

Slow-release Dex implants were prepared to deliver 0.5 mg/g body weight Dex in 100 μL of vegetable shortening/peanut oil and administered as in Phase 1 using a new Dex source (142255-5G, lot 50033374; Beantown Chemical, Hudson, NH, USA). We administered Dex or control implants to each fish (30 per treatment group in respective tanks). Tanks were equally distributed between experimental diets: Bio-Oregon® BioVita Fry 1.2mm pellet diet at 3.5% fish biomass daily (control diet) and food withheld (fasting diet).

Twenty-three days after Dex treatment, fish in the ASE groups were given an inoculum of fresh gastrointestinal tract tissue (mixed mid-intestine and pyloric caeca) from ASE-affected spawning Chinook Salmon as described in section 2.5. Fish were anesthetized and administered 0.5 mL per fish of the inoculum or control (sterile 1x PBS) via gastric gavage using a sterile syringe fitted with a 12-gauge needle and red rubber catheter. At 28 days post-exposure to ASE, eight fish from each tank and treatment group were randomly selected for humane euthanasia and fixed for histology (Figure 1). Additionally, a section of mid-intestine was collected, stored on ice, and frozen at -80°C for future research. Then at 48 days post-exposure to ASE, ten fish from each tank and treatment group were randomly selected for euthanasia and fixation for histology. All remaining fish were euthanized, stored on ice, and frozen whole at -20°C.

Throughout the study period, any fish that became moribund (generally ataxic, loss of buoyancy control, and/or increased respiration rate) were euthanized and are categorized as mortalities for survival analyses.

### 2.5 ASE-affected tissue inoculum

The tissue inoculum used in Phase 2 was collected from adult Chinook Salmon carcasses of Willamette Hatchery broodstock immediately after fish were euthanized and artificially spawned for propagation at South Santiam Hatchery in Sweet Home, Oregon, USA (44°24’57.62”N, 122°40’30.82”W) in September 2023. The tissue inoculum was comprised of fresh gastrointestinal tract tissue (mixed mid-intestine and pyloric caeca) from 20 individual fish and held in plastic bags on ice for six hours before processing. A portion of intestine from each fish was collected in 10% NBF and processed as described in section 2.2 and a subset of 15 fish were screened by histopathology to confirm ASE lesions and presence of *E. schreckii* and *C. shasta*. Tissues were macerated using a stomacher (Colworth 80, Bedfordshire, UK) in equal parts sterile 1x PBS and strained using a 500 µm screen. The homogenate was stored at 4°C for 12 hours prior to administration to experimental fish in section 2.1.2 Phase 2. A subsample of the tissue inoculum was stored at -80°C for future studies.

Control inocula from otherwise healthy spawning spring Chinook salmon (i.e., without ASE lesions) were not available from our sampling population. All fish sampled from the Willamette Hatchery broodstock had ASE lesions confirmed by histopathology. Introducing tissues from a different broodstock population or hatchery would introduce confounding variables difficult to separate in this transmission experiment. Thus, we used sterile 1x PBS as our control inoculum.

### 2.6 Electron microscopy

We examined recipient intestinal tissues from Phase 2 by electron microscopy to investigate the potential presence of viral agents. This analysis was motivated by the discovery of structures suggestive of viral infection in enterocytes of fish with ASE, previously examined by routine transmission electron microscopy (TEM) during the description of *E. schreckii*. Those tissues were from an adult Willamette Hatchery spring Chinook Salmon with ASE, collected in 2019 (Couch et al., 2022). The TEM image of virus infected cells from the previous study (Couch et al., 2022) used FEI Tecnai 12 TEM (FEI, Hillsboro, OR) and the image was recorded with a OneView 16 Megapixel camera using Digital Micrograph software (Gatan, Pleasanton, CA) at the Rutgers Electron Microscopy Facility (Newark, NJ).

Intestines of the one fish from the DexFast + ASE group was processed by correlative light electron microscopy (CLEM), a technique that enables the examination of the same structures observed in histological slides by TEM (Dobbie, 2019). Histologic regions of interest were identified by observing HCE-stained sections and matched to corresponding regions on slides of unstained paraffin sections. The areas of interest on unstained slides were marked using a diamond scribe. Slides from one of the recipient fish with intranuclear inclusions in enterocytes were subsequently deparaffinized, rehydrated, and processed for CLEM as follows. The sample was fixed with 2.5% glutaraldehyde, 2.0% paraformaldehyde in 0.1 M sodium cacodylate buffer, post fixed with 1% osmium tetroxide in buffer, and *en bloc* stained with 2% aqueous uranyl acetate, dehydrated through a graded series of ethanol, and embedded in Lx112 resin using an inverted BEEM capsule, with its truncated tip removed, over the area of interest. After polymerization the slide was gently heated and the BEEM capsule containing the embedded tissue was peeled off the slide. Using gentle heating, the Epon embedded tissue block was removed from the capsule and trimmed. Ultrathin (70 nm) sections of the scribed area were cut on a Leica EM Ultracut UC7, stained with uranyl acetate followed by lead citrate. CLEM sections were viewed on a JEOL 1400 transmission electron microscope (Peabody, MA) and images were collected with a Gatan Orius camera using Digital Micrograph software (Gatan, Pleasanton, CA). CLEM processing and imaging was conducted at the Albert Einstein College of Medicine Analytical Imaging Facility.

### 2.7 Statistical analysis

All statistical analyses were performed using R (R Core Team, 2024) and RStudio (Posit team, 2024). For Phase 1 mortality data, days 19 to 25 were excluded from the analysis to remove the period of the disease outbreak.

Kaplan-Meier curves were constructed using the *survival* package version 3.6-4 (Therneau, T, 2024; Therneau & Grambsch, 2000) and the *ggsurvfit* package version 1.1.0 (Sjoberg et al., 2024) to visualize survival probabilities over time. For Phase 1, the survival curves were stratified by Dex treatment and diet treatment. For Phase 2, the survival curves were stratified by the combination Dex and diet treatment and the ASE exposure treatment with the following treatment groups: ASE exposure (ASE Only); control (Control); Dex, fasting, and ASE exposure (DexFast + ASE); or Dex and fasting (DexFast Only).

Cox Proportional Hazard models were fit using the *survival* package to assess the effects of treatment variables on survival in both Phase 1 and Phase 2 experiments. To quantify the magnitude of the treatment effects, Cohen’s *d* was calculated as a measure of effect size. This provided a complementary metric to the hazard ratios from the Cox models. Hazard ratios (HR) were calculated as the exponentiated coefficients from the Cox models, representing the relative risk of mortality between treatment groups and controls. Binary indicator variables (0,1) were created for treatment variables, Phase 1 Dex and diet treatments and Phase 2 ASE exposure and combination Dex and diet treatment, with the respective control as the statistical baseline. The initial Phase 1 model was evaluated for proportional hazards assumptions using Schoenfeld residuals (Grambsch & Therneau, 1994), revealing a violation for the low dose Dex treatment. To address this, the model was adjusted to allow the effect of low dose Dex treatment to vary over time using the tt() function in the *survival* package. This adjustment accounted for the non-proportionality observed in the initial analysis. For Phase 2, the initial model included an interaction term between Dex-fasting and ASE treatments. We compared this model with a simpler model without the interaction term using Akaike’s Information Criterion (AIC). The proportional hazards assumption was tested using Schoenfeld residuals, revealing violations for both Dex-fasting and ASE treatments. To address this, the model was adjusted to allow the effects of Dex-fasting and ASE treatments to vary over time.

Although the histopathologic assessments were scored on ordinal scales, the differences in the severity of histopathologic lesion of adjacent categories were approximately equal (e.g., A was 50% of B and B was 50% of C). Therefore, we used an ANOVA to assess overall differences in various variables across treatment groups. In Phase 1, these variables included enteritis, intestinal folds, hepatic vacuolization (lipidosis and glycogenosis), kidney melanin, coelomitis, branchitis, and myositis. In Phase 2, the variables were enteritis, weight, intestinal fold loss, hepatic vacuolization (lipidosis and glycogenosis), kidney melanin, coelomitis, myositis, and intestinal epithelial dysplasia. Differences were considered statistically significant when p < 0.05. For significant ANOVA results, Tukey’s Honest Significant Difference (HSD) tests were performed for post-hoc pairwise comparisons to identify specific differences between treatment groups.

Violin plots were generated using *ggplot2* package (Wickham, 2016) to display the distribution of histology scores across the four treatment groups.

## 3 Results

### 3.1 Phase 1

#### 3.1.1 Survival Analysis

The Kaplan-Meier curves revealed distinct survival patterns across different treatment groups (Dex dose and diet) in the Phase 1 experiment (Figure 2). We excluded days 19 to 25 to ensure that the analysis focused on the treatment effects without the confounding influence of the *I. necator* and *Saprolegnia* spp. epizootic diagnosed tank-side by wet mount and macroscopic examination. The infections were confirmed with a subsampling on day 25, where moribund and a random sampling of healthy fish were selected for histopathology from high fast tanks (A and B) and control tank E. Sampled fish included high fast and high dose Dex (n=7 total, 1 moribund), high fast and low dose Dex (n=2 total, 1 moribund), and controls (n=3).

**Figure 2.**
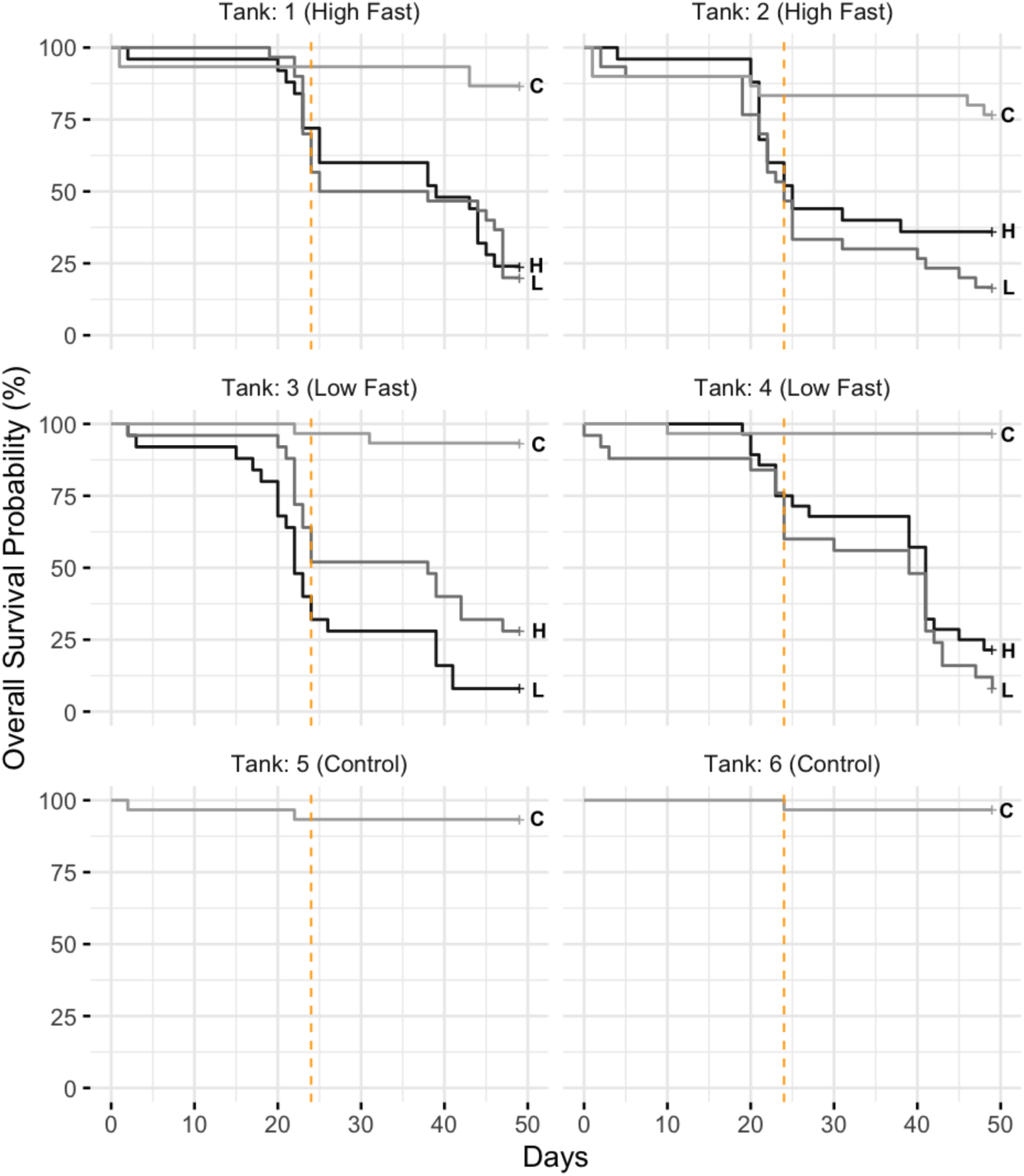
**Phase 1: Kaplan-Meier survival curves**, where the high dose (H) and low dose (L) dexamethasone (Dex) treatments significantly increased the risk of mortality (p < 2 X 10^-16^) compared to controls (C) with the high dose (H) Dex treatment having the strongest effect. The low dose treatment also varied over time and was modeled as a time-varying coefficient. A Cox proportional hazards model was used to assess treatment and tank effects. The two overlapping treatment groups (in duplicate) are 1) diet treatment including normal feed (control; Tanks 5 and 6), low fast (Tanks 3 and 4), and high fast (Tanks 1 and 2). The vertical dashed line indicates when a formalin bath treatment was administered for an *Ichthyobodo necator* infection at 24 days post-Dex treatment.

The survival curves, stratified by Dex treatment (Control, Low Dose, High Dose) and diet (Control, High Fast, Low Fast), showed varying survival probabilities over time. Visual inspection of the curves indicated that the Dex treatment had a more pronounced effect on survival compared to the diet treatments. Minimal fish mortalities occurred in fasting only groups, in contrast to high mortalities and variations in survival probabilities that were observed within Dex treatment groups.

The adjusted model identified significant effects for both low dose and high dose Dex treatments on survival. Specifically, 1) high dose Dex treatment was associated with a significantly increased risk of mortality, with a very large effect size: hazard ratio (HR) = 7.3, 95% CI: 4.7-11.3), representing a 7.3-fold increase in mortality risk compared to the control group (*p* < 2x10^-16^), 2) the low dose Dex treatment also showed a significant cumulative effect on mortality (HR = 1.3, 95% CI: 1.2-1.4, p <2x10^-16^), but this effect varied over time, and 3) the effects of high fast (HR = 3.5, 95% CI: 1.0-11.6, *p* = 0.044) and low-fast diet treatments (HR = 3.5, 95% CI: 1.1-11.8, *p* = 0.041) were marginally statistically significant in the final model. However, it still represented a substantial effect size with both diet treatments associated with approximately 3.5-fold increased risk of mortality. No difference was seen between Control subgroups – i.e., those receiving the sham implants and those that did not in Tanks 5 and 6 (Figure 2).

#### 3.1.2 Phase 1 Histopathology

The following changes were observed across the different groups: branchitis, enteritis, intestinal fold reduction, changes in enterocyte height, vacuolation of hepatocytes, increased kidney melanin deposition relating to diet and coelomitis scores were based on a range of 0-3. Renal melanosis was the only pathologic change that was statistically significant between groups.

##### Gills

Severe branchitis was only observed in fish treated with Dex early in the experiment, including three moribund (High Fast + High Dex, n=1; High Fast + Low Dex, n=2) and six non-moribund fish (High Fast + High Dex, n=6). Microscopic gill disease noted by histopathology (Figure 3) included *I. necator* (n=2, high fast and high Dex dose), consistent with the wet mount observations, numerous suboval or pyriform protozoa identified as *I. necator* were observed (Figure 3). Two fish at the termination of the experiment demonstrated mild, mononuclear branchitis without apparent pathogens (High Fast Only, n=1; Low Fast + Low Dex, n=1), which was not statistically significant. No pathologic changes were observed by histology in control fish (n=3).

**Figure 3.**
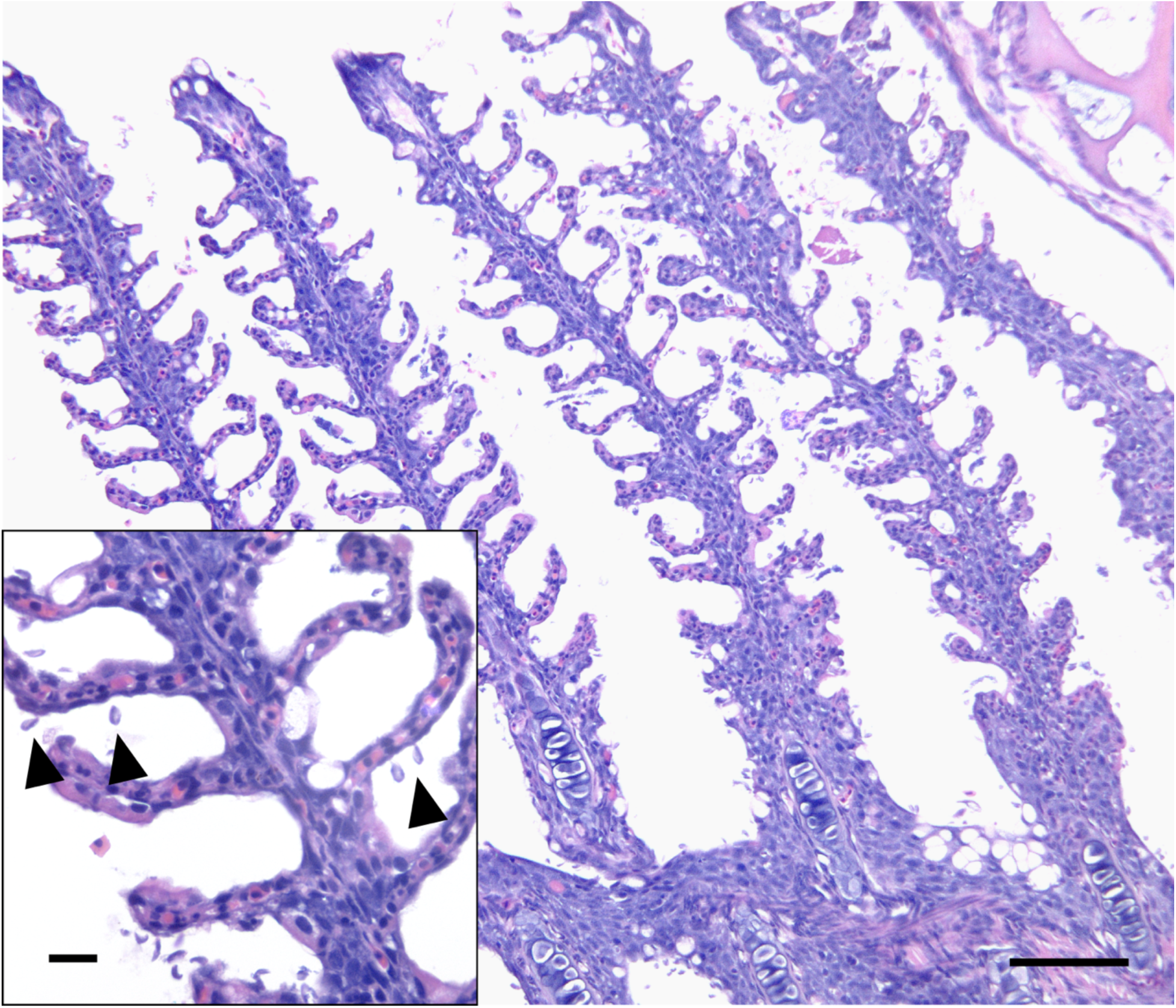
Phase 1 Gill. Representative histology of branchitis. Moderate inflammatory infiltrate expands the interstitium of the primary and secondary filaments, comprised of lymphocytes and granulocytes. The secondary filaments are blunted with epithelial hyperplasia and fusion. At higher power (inset), large numbers of *Ichthyobodo necator* (arrowheads) are attached to the gill epithelium. HCE stain. Scale bar = 100 µm, inset 20 µm.

##### Intestine

Decreased enterocyte height and cytoplasmic volume, as expected with fasting and Dex, was appreciated by histopathology (Figure 4). Whereas not subjected to statistics, fasted fish appeared to have reduced height of enterocytes and a mild reduction in the appreciated numbers of goblet cells in the mid-intestinal sections. No statistically significant differences in intestinal fold heights or enteritis scores were observed, supported by all fish in all treatment groups that were examined (n=31) having normal intestinal folds (score 0) and one fish demonstrating mild enteritis (score 1).

**Figure 4.**
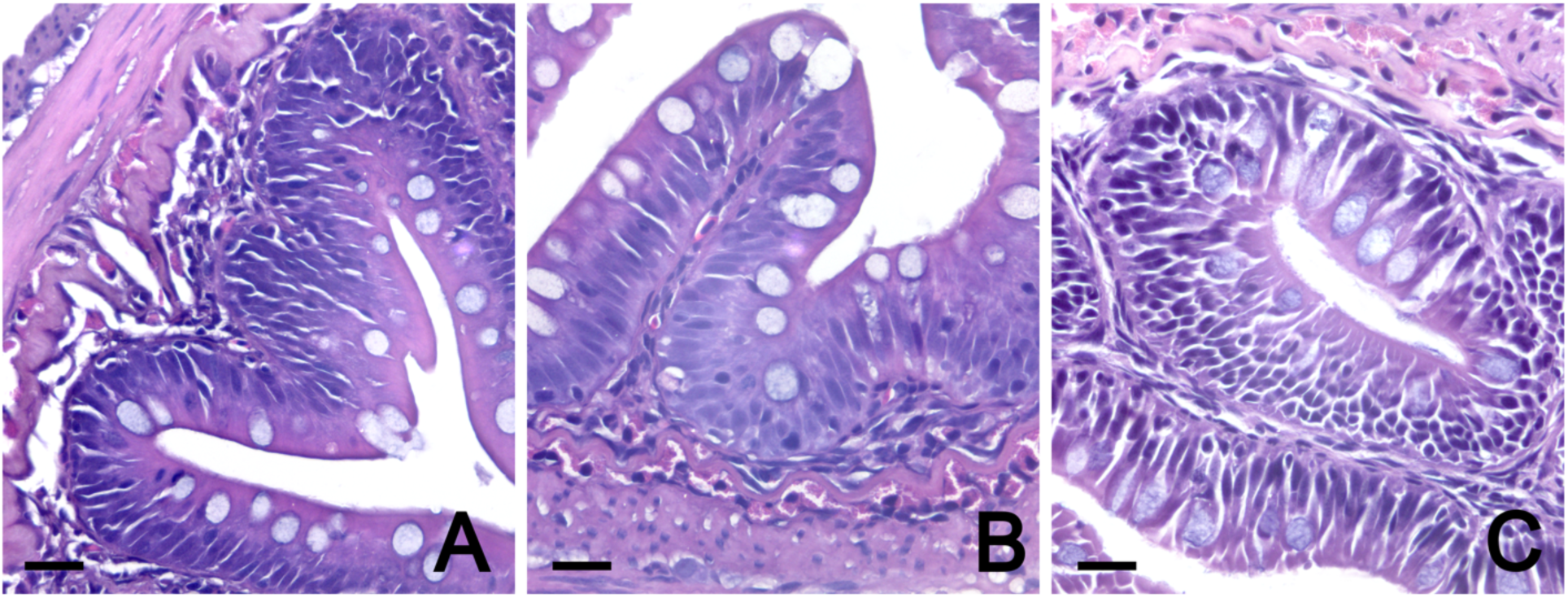
Phase 1 Mid-intestine. Enterocyte morphology with dexamethasone (Dex) and fasting treatment; A) control, B) high dose Dex, and C) high fast fish. Fish under a fasting regime demonstrate decreased cytoplasmic volume of the enterocytes (width and height). HCE stain. Scale bar = 20 µm.

##### Kidney

The deposition of melanin within the renal interstitium (Figure 5, 6) coincided with diet regimens. Compared to the Control group with sham implants, the High Fast + High Dex group (*p* = 0.0025) and the High Fast Only (*p* = 0.010) group had significantly higher renal interstitial melanin deposition (Figure 6). These data included both 24-days post-Dex exposure for the High Fast + High Dex group due to no fish in that group surviving until 50-days post-Dex exposure, while all other groups had fish that survived until this later time. Overall, these analyses showed that reduced feed (particularly fasting) was positively associated with increased melanin deposition in the renal interstitium.

**Figure 5.**
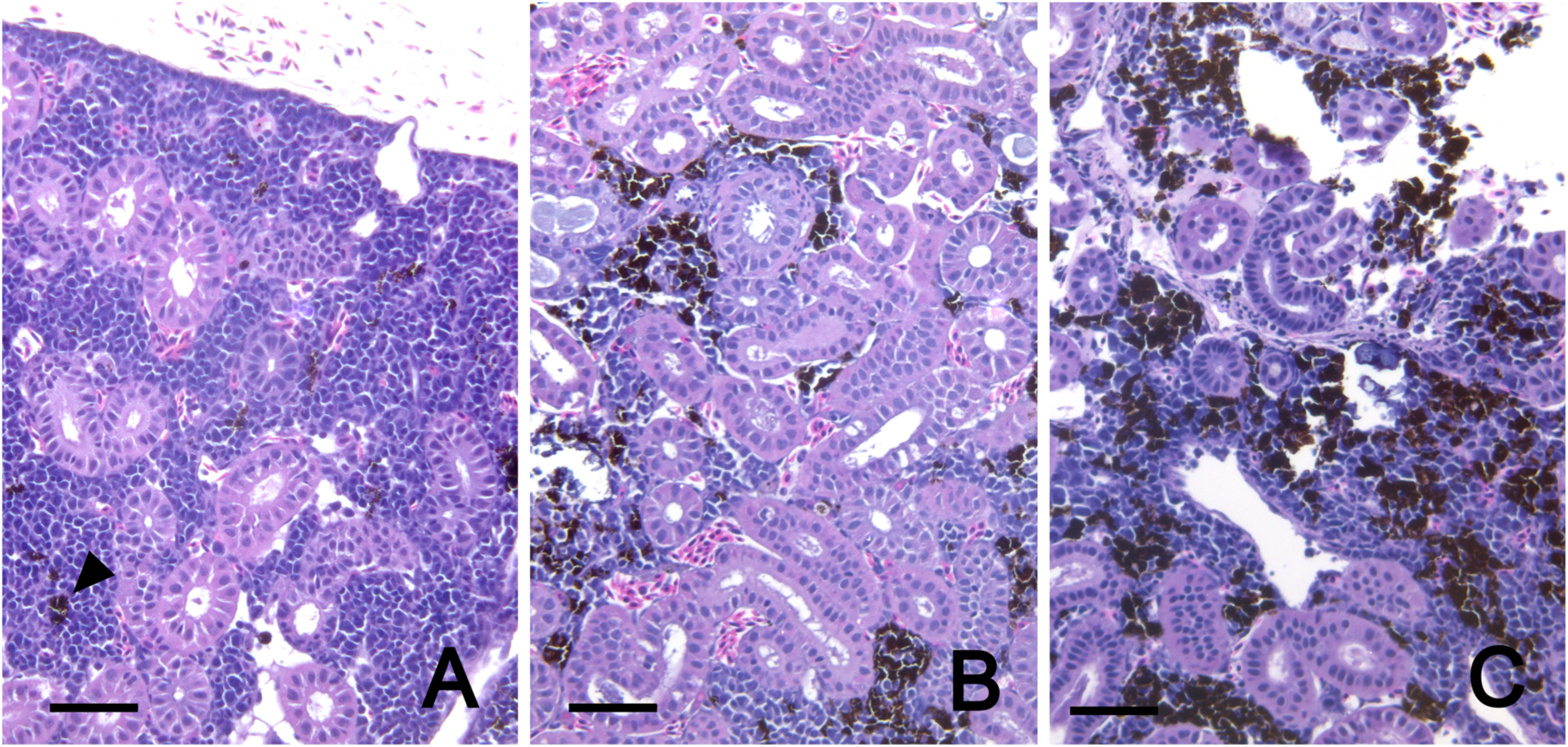
Phase 1 Kidney. Renal melanosis scores with representative histology based on abundance of dark brown to black granular pigment in the intersitium. A) Mild melanin deposition with occasional small clusters of melanin (arrowhead) = score 1, B) moderate melanin deposition = score 2, and C) severe melanin deposition = score 3. HCE stain. Scale bar = 50 µm.

**Figure 6.**
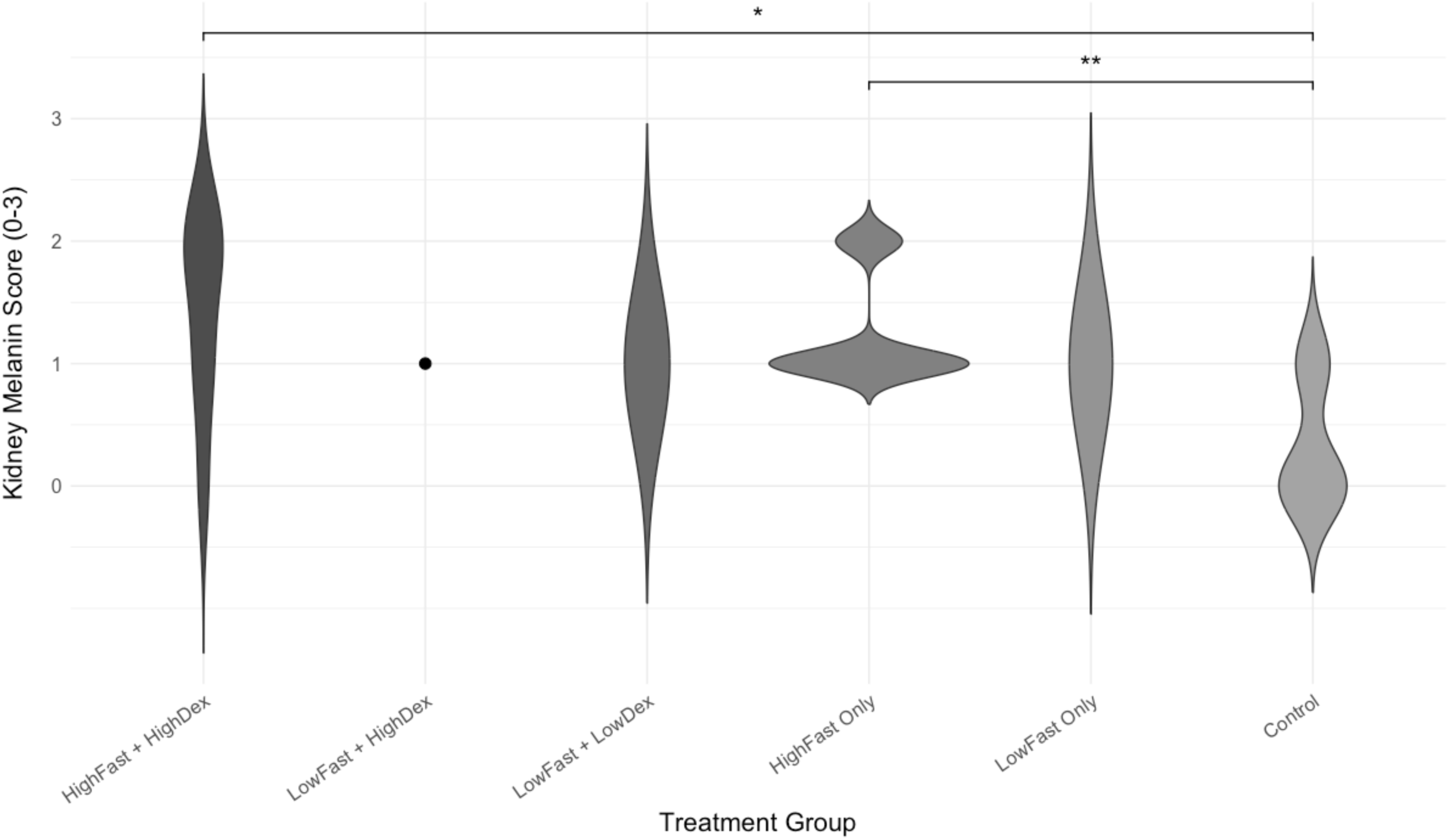
Phase 1: Violin plots for renal melanosis. High fasting treatment significantly increases interstitial melanosis compared to the control group, regardless of dexamethasone treatment based on histopathology results. There are six treatment groups shown: HighFast + HighDex, LowFast + HighDex, LowFast + LowDex, HighFast Only, LowFast Only, and Control, where dexamethasone and a fasting diet simulate immunosuppression and anorexia of adult spawning salmon. Fish in the HighFast + HighDex group were sampled at 24 days post-exposure as very few in this group survived to the 50-day post-exposure termination of the experiment. The kidney melanin scores range from 0 to 3, where 0 is no melanin in the renal intersitium, 1 is mild, 2 is moderate, and 3 is prominent (see Figure 4). Tukey HSD Test. An asterisk (*) between the groups indicated statistical significance, where <0.001 = ** and <0.01 = *.

##### Liver

Livers from all fish across the groups where within normal limits regarding hepatocyte vacuolar change (Figure 7), and no statistical differences between this endpoint were observed (See Supplemental Data).

**Figure 7.**
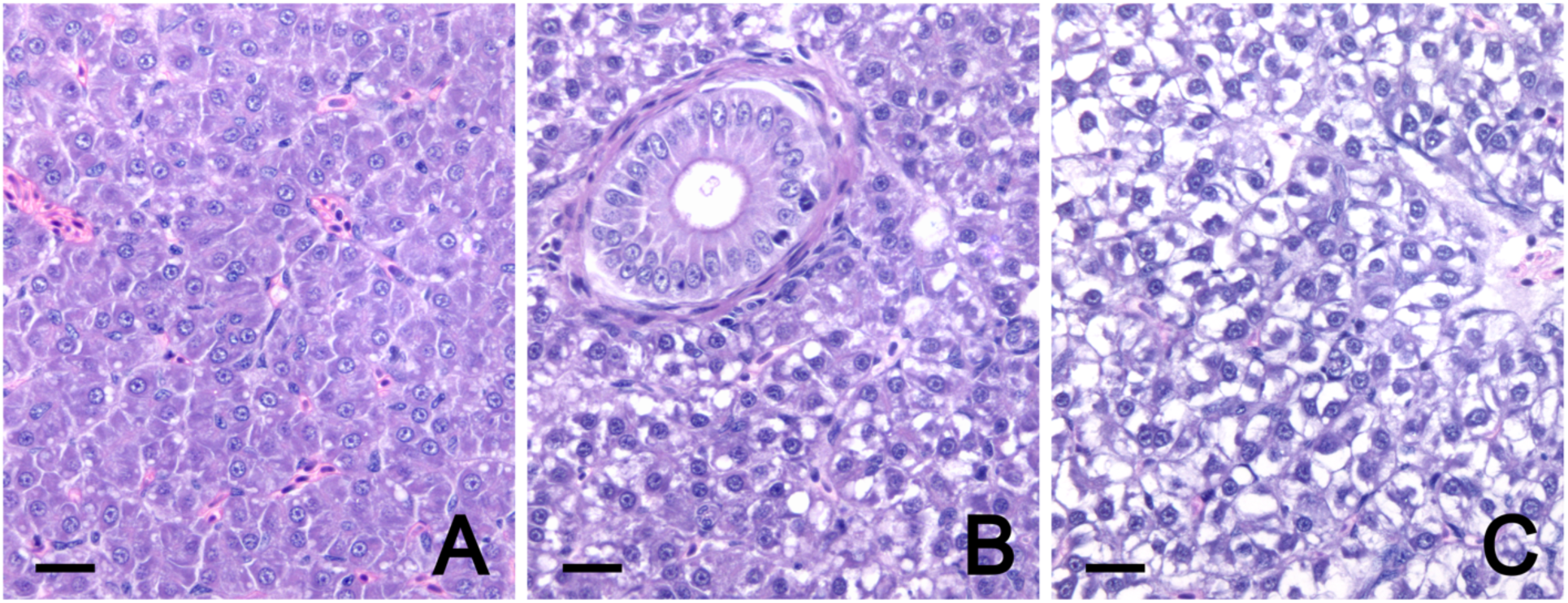
Phase 1 Liver. Representative histology of vacuolar change scores in hepatocytes based on severity. A) mild change = score 1 (Control), B) moderate change = score 2 (Control), and C) severe change = score 3 (Control). HCE stain. Scale bar = 20 µm.

##### Coelom

Inflammation of the coelomic space, coelomitis, was observed in a total of five fish: one Control, two High Fast Only, one Low Fast Only, and one Low Fast + Low Dex fish. Histologic findings of coelomitis included focally extensive regions of fibroblastic proliferation associated with the surface of the visceral adipose and imbedded pancreatic tissues with mild to marked numbers of percolating foamy macrophages and mononuclear leukocytes. The foamy to large cytoplasmic vacuoles in the macrophage population was indicative of a response to the free lipid within the IP vegetable shortening or peanut oil vehicle used for Dex and sham treatment.

### 3.2 Phase 2

#### 3.2.1 ASE-affected tissue inoculum

For the 20 fish used to make the inoculum, ASE-lesions were observed in all fish examined by histology (n=15) with a mean of 60% epithelium remaining. *E. schreckii* was present in 5 of the 15 histology samples, ranging from mild to moderate infections (Nervino et al., 2024). *C. shasta* was present in all fish examined, and three fish contained luminal larval cestodes.

#### 3.2.2 Survival

The Kaplan-Meier survival analysis for the Phase 2 experiment revealed distinct survival patterns across the different treatment groups (Figure 8). The survival curves, stratified by treatment, indicated that the DexFast + ASE exposed fish across replicate tanks exhibited the lowest survival probability over time. This was followed by the DexFast Only (without ASE). Indeed, no fish in the DexFast + ASE survived past 50 days post exposure, and only 30% of the fish in the DexFast without ASE survived until the end of the experiment. In contrast, the other groups that were not Dex treated or fasted, including the group exposed to ASE, showed minimal mortalities (Figure 8).

**Figure 8.**
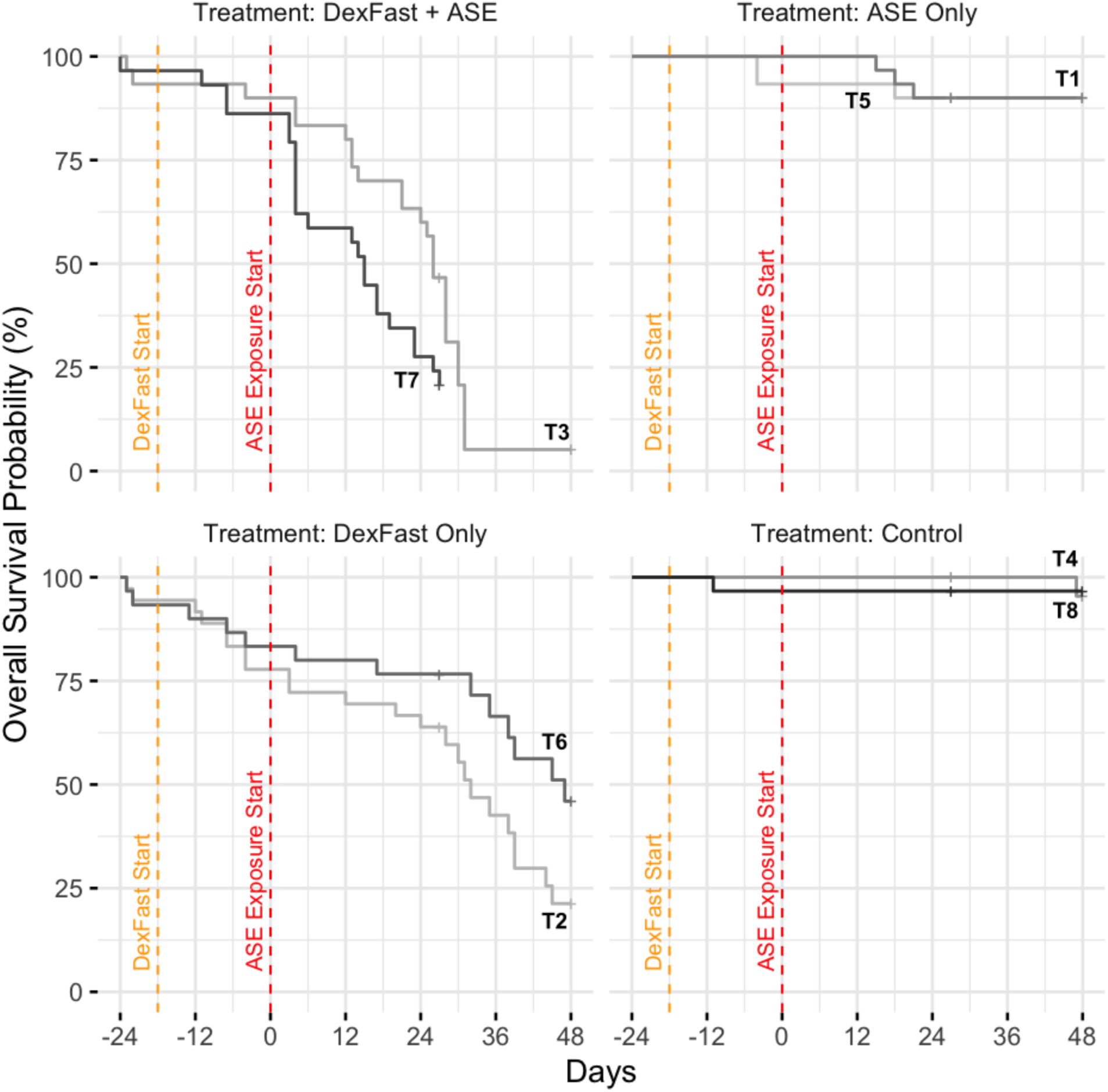
**Phase 2: Kaplan-Meier survival curves** by treatment and duplicate tanks. Treatment groups include DexFast and ASE exposure (DexFast + ASE), ASE-exposure (ASE Only), DexFast Only, and Control, with each group divided between two tanks (T). Fasting diet was the same as High Fast in Phase 1 – i.e., no feed). Both DexFast and ASE treatments individually increased mortality risk over time (p < 0.001 for both treatments), with DexFast having a much stronger effect, based on a Cox proportional hazards model. The DexFast treatment increased the hazard of mortality 2-fold, where the ASE treatment increased the hazard of mortality 1.5-fold.

The Cox Proportional Hazards model further verified the negative impact of both Dex-Fast and ASE treatments on risk of mortality, with the DexFast treatment having a stronger effect. The initial model, which included an interaction term between DexFast and ASE treatments, indicated a significant effect of these treatments on survival. However, the model without the interaction term provided a better fit, as evidenced by a delta AIC of 5.36. The effects of DexFast and ASE treatments on fish mortality were evaluated using a time-varying Cox proportional hazards model. The model demonstrated a significant improvement in fit compared to traditional Cox models, as indicated by a delta AIC of 3.41. The time-varying Cox model showed that DexFast treatment significantly increased mortality risk (HR = 2.08, 95% CI: 1.72-2.50, p < 0.001), with the magnitude of this effect increasing over the course of the experiment. Similarly, the time-varying Cox model showed that ASE treatment significantly increased mortality risk (HR = 1.45, 95% CI: 1.25-1.68, *p* < 0.001), with this effect also increasing over time, though to a lesser extent than DexFast. Overall, the analyses confirmed that DexFast and ASE exposure significantly increased risk of mortality.

#### 3.2.3 Phase 2 Histopathology

Differences in pathologic changes were observed microscopically between the treatment groups, which is summarized in Table 3. Changes assessed included enteritis, intestinal fold loss, hepatic vacuolization (lipidosis or glycogenosis), interstitial kidney melanin, coelomitis, intestinal epithelial dysplasia, and myositis. Statistical analysis revealed a significant ANOVA result for enteritis, kidney melanin, intestinal fold loss, and intestinal epithelial dysplasia. Intestinal fold loss had no significant difference between treatment groups after a Tukey HSD test analysis and was removed from the variables of interest. Intestinal fold loss, hepatic vacuolization, coelomitis, and myositis, were not statistically significant (*p* > 0.05).

**Table 3.**
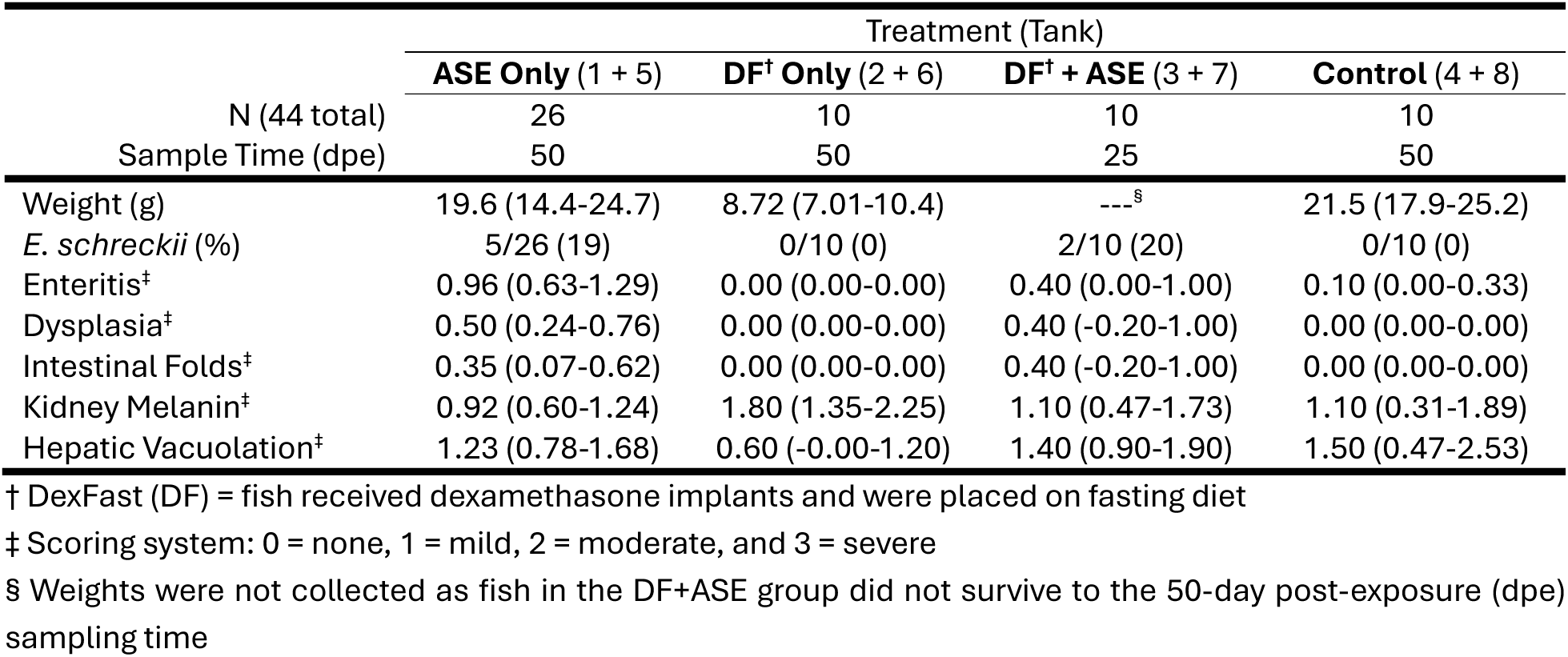
**Phase 2: Histopathologic findings** in Chinook Salmon exposed to Adult Salmon Enteritis (ASE) tissues with or without immunosuppression (dexamethasone implant and fasting). Control fish received neither DexFast^†^ nor ASE inoculum. Pathology scores are mean (95% confidence interval) and *Enterocytozoon schreckii* is prevalence based on results from HCE or Luna-stained slides combined.

##### Intestine

Enteritis occurred within the pyloric caeca and mid-intestine and was primarily composed of small mononuclear cells, consistent with lymphocytes and some plasma cells, with rare granulocytes within the lamina propria (Figure 9). The inflammatory infiltrate would periodically expand into the overlying epithelium as individual cells or small clusters. Throughout the affected intestines, there were foci of epithelial dysplasia characterized by cytomegaly, rounding, piling or disorganization of cells, cytoplasmic vacuolar change, and loss of polarity. Large, basophilic to amphophilic, sometimes glassy, intranuclear inclusion bodies were noted within the enterocytes of two fish with enteritis and dysplasia (Figure 10B, E), one DexFast + ASE and one ASE Only. To further characterize these putative viral inclusions, CLEM was performed on formalin-fixed paraffin wax embedded tissues from a target area of the mid-intestine from the former (Section 3.3 Electron Microscopy). The Tukey HSD test demonstrated significant increase enteritis scores in the ASE Only group compared to all other treatment groups (Figure 11). The significant differences between groups are indicated by an asterisk in Figure 11, where ASE Only had significantly higher enteritis scores compared to Control (*p* = 0.003), DexFast Only (*p* = 0.001), and DexFast + ASE (*p* = 0.04) groups.

**Figure 9.**
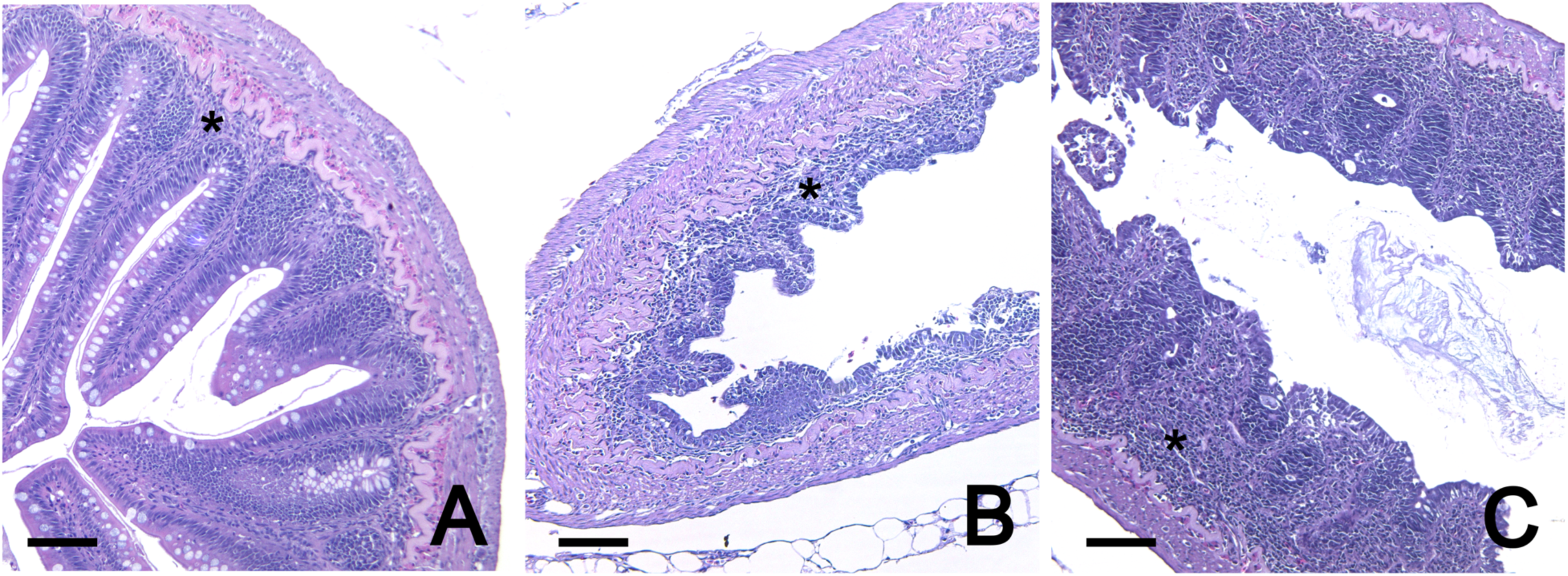
Phase 2 Mid-intestine. Histologic sections of Chinook Salmon mid-intestine showing representative enteritis scores and intestinal fold changes. Enteritis was scored on a scale of 0-3 (0 = normal/no inflammation, 1 = mild, 2 = moderate, 3 = severe), and intestinal fold height changes were scored on a scale of 0-3 (0 = normal height, 1 = mild reduction, 2 = moderate reduction, 3 = severe reduction). A) Control sample with mild inflammation (score 1) in the lamina propria (asterisk) and normal intestinal fold height (score 0). B) Moderate enteritis (score 2) with moderate reduction in intestinal fold height (score 2) and expansion of inflammation of the lamina propria (asterisk) in a DexFast + ASE treated fish. C) Severe enteritis (score 3) with moderate reduction in intestinal fold height (score 2) and severe expansion of inflammation of the lamina propria (asterisk) in ASE only treated fish. HCE stain. Scale bar = 100 µm.

**Figure 10.**
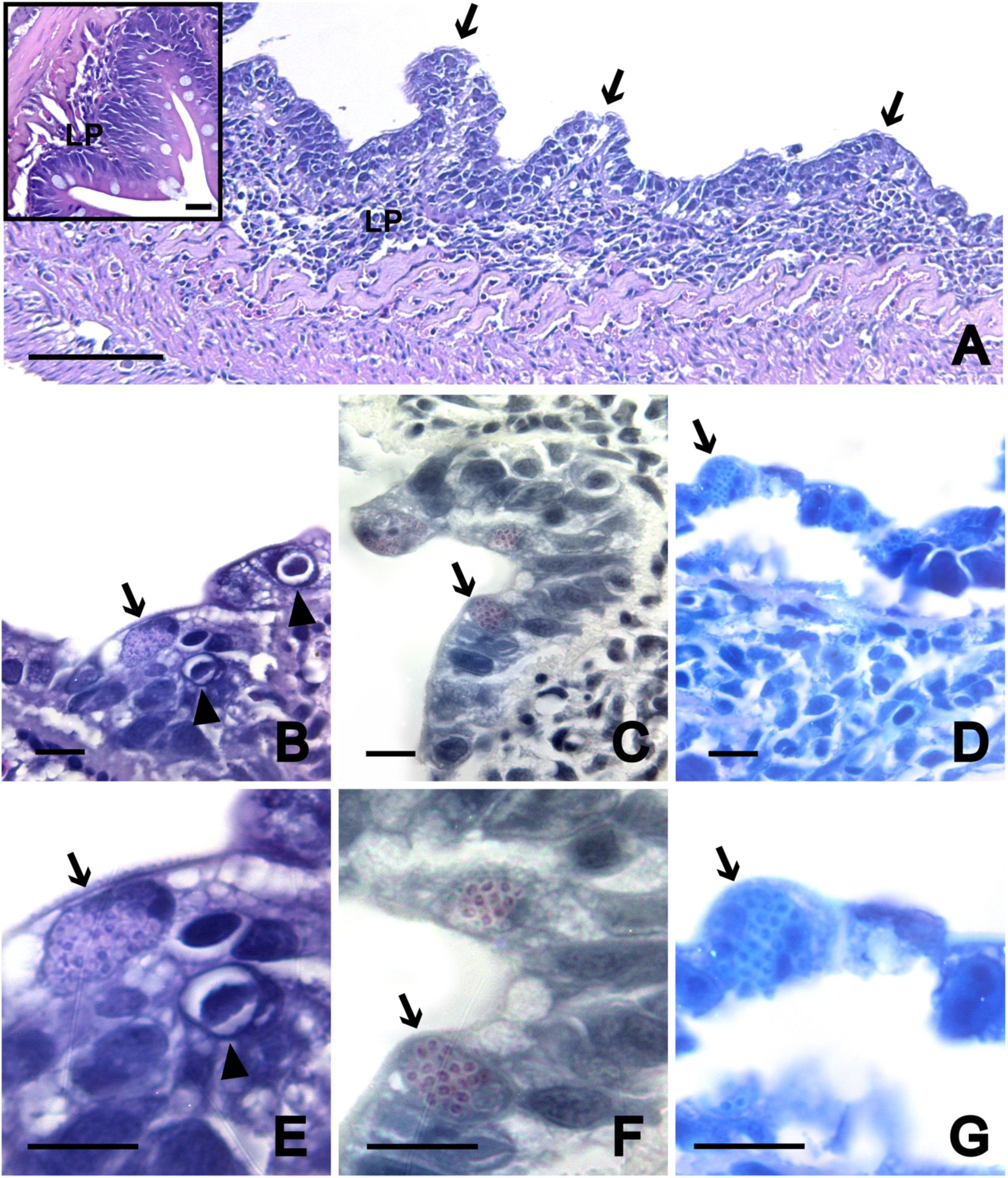
Phase 2 Mid-intestine. Histologic sections of Chinook Salmon intestinal epithelium showing *Enterocytozoon schreckii* infection and putative viral inclusions in enterocytes of a DexFast + ASE treated fish. A) Section of mid-intestine from mucosal to serosal surface demonstrating a moderate enteritis (score 2) comprised of lymphocytes, plasma cells, and rare granulocytes predominately expanding the lamina propria (LP) with a few inflammatory cells percolating into the mucosal epithelium. The intestinal folds are reduced (score 2) with epithelial attenuation and enterocyte apical shortening, cytoplasmic basophilia and vacuolation, and dysplasia (score 2). Arrows indicate areas targeted for correlative light electron microscopy (CLEM) in Figure 15. Mid-intestine from a control group, no Dex or fasting, is shown as an inset. HCE stain. Scale bar = 100 µm; inset = 20 µm. B-E) Higher magnification of the intestinal mucosa shown in A, where aggregates of *E. schreckii* spores (arrows) and intranuclear virus-like inclusions (arrowheads) are indicated. B, E) HCE stain. Scale bar = 10 µm. C, F) Luna stain. Scale bar = 10 µm. D, G) Giemsa stain. Scale bar = 10 µm.

**Figure 11.**
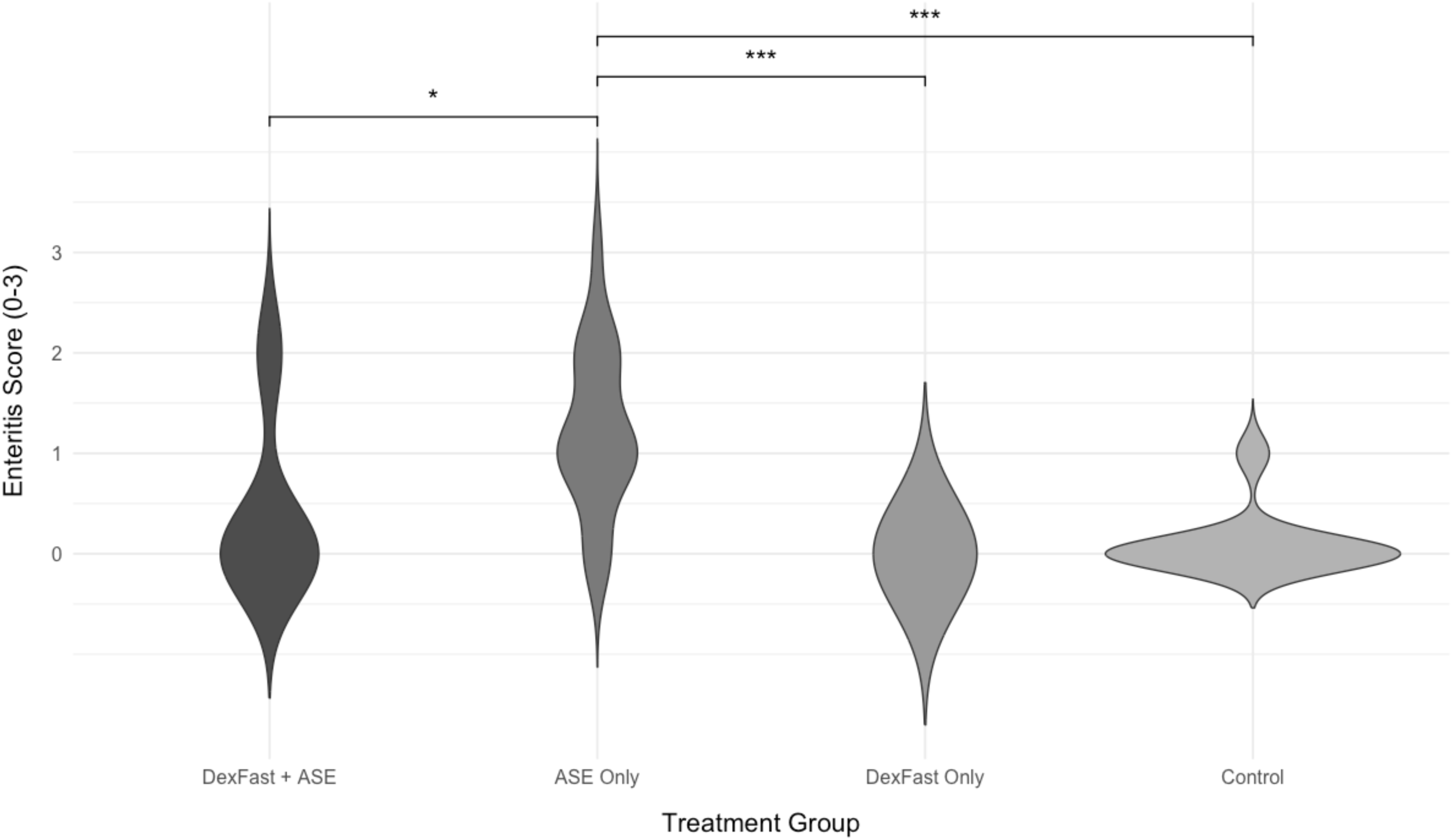
Phase 2 Enteritis. Four treatment groups: Control, DexFast Only (treatment with dexamethasone and a fasting diet), ASE Only (exposure to the ASE tissue inoculum), and DexFast + ASE. Fish exposed to ASE had significantly increased enteritis. Enteritis was scored on a scale of 0-3 (0 = normal/no inflammation, 1 = mild, 2 = moderate, 3 = severe). Tukey HSD Test. An asterisk (*) between the groups indicated statistical significance, where <0.0001 = *** and <0.01 = *.

The DexFast Only group exhibited a lower median and a narrower distribution of enteritis scores (Figure 11), which was visually represented by the histopathologic enteritis results (Figure 9B), suggesting the DexFast treatment may have had a protective effect against enteritis. Likewise, the protective effects of the DexFast treatment on enteritis scores are similarly visualized between the DexFast + ASE group and the ASE Only group (Figure 11). In contrast, the ASE Only group showed a wider distribution and higher median scores, indicating a potential exacerbation of enteritis with ASE exposure (Figure 9C). The Control, DexFast Only, and DexFast + ASE groups displayed intermediate distributions, with no significant differences observed between them.

Dysplastic change of the intestinal epithelium, a hallmark of ASE in fish from the field (Nervino et al., 2024), was significantly increased in the ASE Only group compared to the Control (*p* = 0.005, *d* = 3.77) and DexFast Only (*p* = 0.005, *d* = 3.77) groups (Figure 12). Some fish within the DexFast + ASE group showed dysplasia, but as only 10 fish were evaluated from this group, with low statistical power, this did not show any statistical significance compared to other groups. Importantly, dysplasia was only seen in groups exposed to ASE tissue – i.e., both the Control and DexFast Only groups had no intestinal dysplasia (scores of 0).

**Figure 12.**
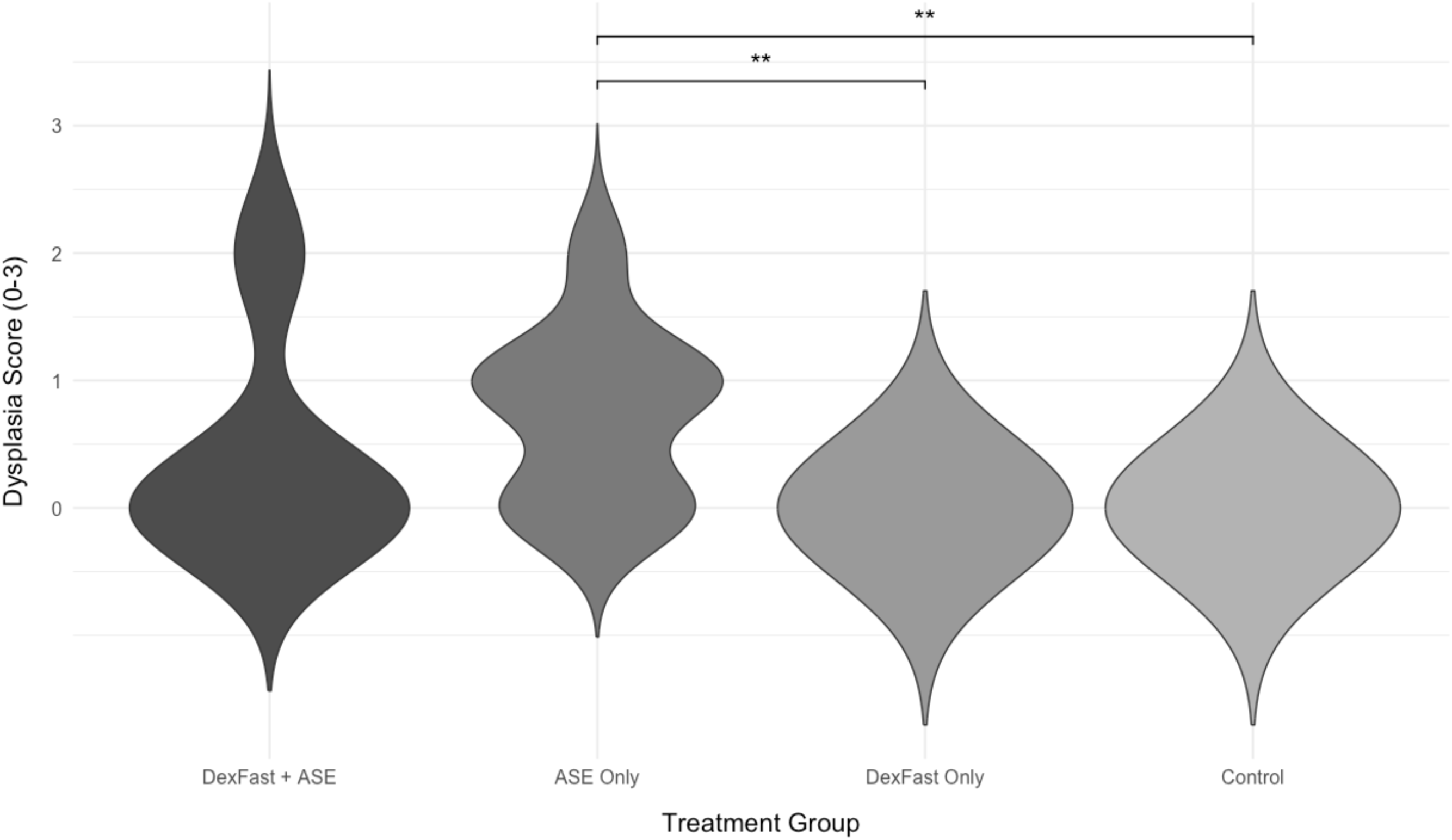
Phase 2 Enterocyte Dysplasia. Dysplasia of the intestinal epithelium was present only in fish exposed to ASE, including both DexFast and ASE Only fish (immunocompetent). Difference relating to ASE exposure was statistically significant between the ASE Only group compared to DexFast Only (*p* = 0.012) and Control (*p* = 0.012) groups. Tukey HSD Test. An asterisk (*) between the groups indicated statistical significance, where <0.001 = **.

##### Kidney

Melanin scores for the renal interstitium were significantly higher in the DexFast Only group compared to the ASE Only group (p = 0.016, *d* = 4.80) using the Tukey HSD test (Table 3). The fasting effect on the DexFast treatment is further demonstrated by the DexFast Only group weighing significantly less than the Control (p = 0.0008, *d* = 8.80) and ASE Only (p = 0.01, *d* = 4.74) groups. We were unable to compare to the DexFast + ASE group, as these fish reached 100% mortality mid-experiment.

##### Parasites

*E. schreckii* was observed in the intestinal epithelium of fish from both DexFast+ASE and ASE Only fish (Table 3), but in no fish from the groups not exposed to the ASE inoculum. Using HCE and Luna stain, *E. schreckii* (Figure 10B,E and C,F, respectively) was detected in the epithelium of 2 of the 10 DexFast + ASE fish, which were sampled at 24-days post-Dex exposure as none survived to the 50-days post-Dex exposure sampling time. Intestinal sections from one fish from the DexFast + ASE was also Giemsa-stained (Table 3, Figure 10D,G), a diagnostic microsporidial stain, which highlighted the microsporidium in enterocytes. The microsporidium was also observed in 5 of the 26 ASE Only fish (Table 3), which were sampled at 50-days post-Dex exposure. The *E. schreckii*-positive fish (n=7) had a mean enteritis score of 1.7, while the intestines of those in which the parasite was not detected but were exposed to ASE inoculum (n=23) had a mean score of 0.67, representing a very large effect size (*d* = 10.3). Fish that were positive for *E. schreckii* had a mean intestinal dysplasia score of 1.6, while those exposed to the ASE inoculum without detectable *E. schreckii* had a mean score of 0.14, also showing a very large effect size (*d* = 14.6).

##### Skeletal muscle

Myositis was observed in a total of two fish in the DexFast Only treatment group (Figure 13), which was not statistically significant. The lesion was comprised of multifocal areas of monophasic, myofiber degeneration in the epaxial and hypaxial muscles accompanied by a mild leukocytic infiltrate.

**Figure 13.**
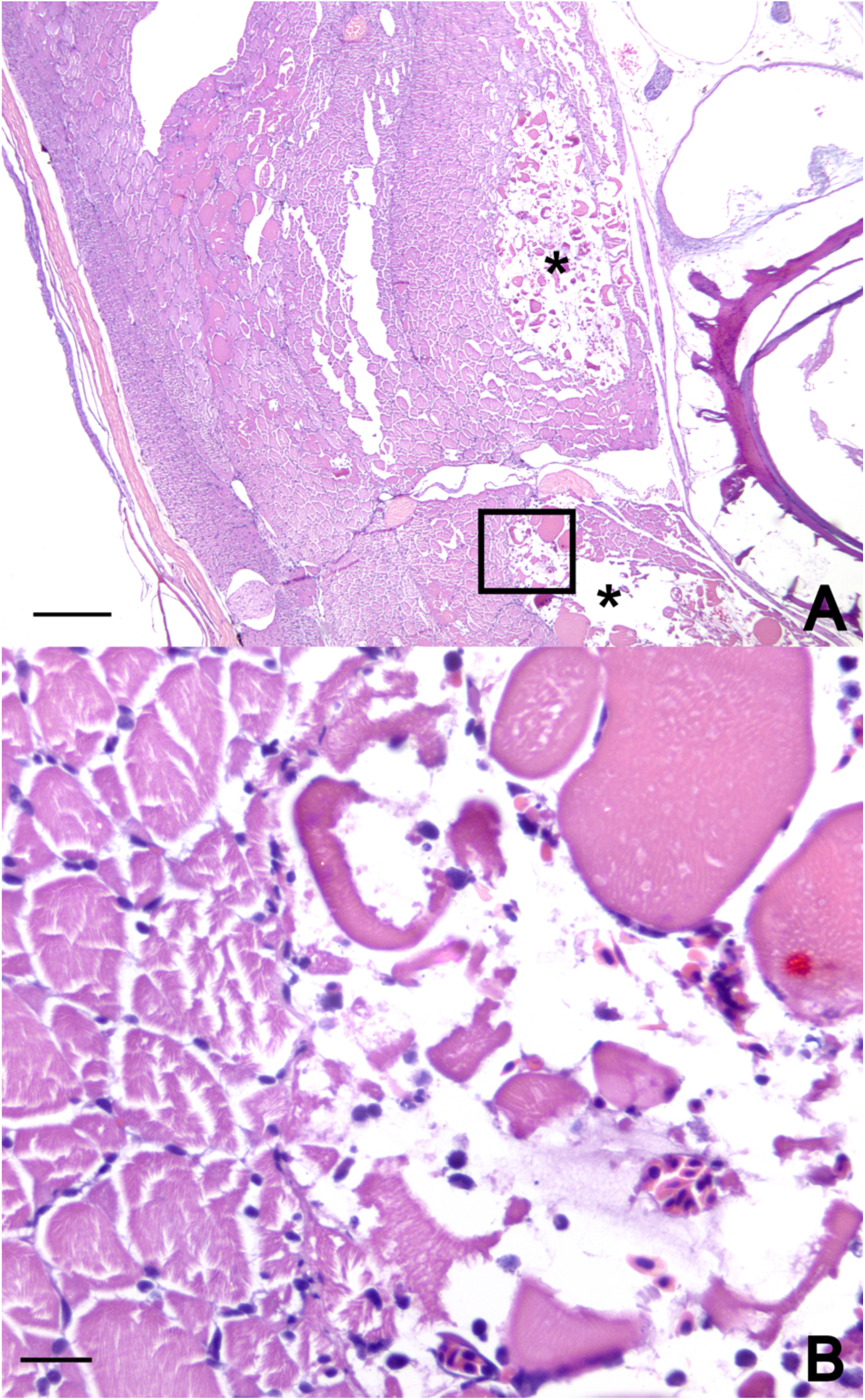
Phase 2 Skeletal Muscle. Representative histology of myositis. A) There are multiple focal areas of myofiber degeneration in the epaxial and hypaxial muscles (asterisks), with one section indicated by a black box for higher magnification. HCE stain. Scale bar = 20 µm. B) At higher magnification, the myocytes in the top right are swollen, hypereosinophilic, with some loss of cytoplasmic striations, the central myocytes are shrunken, hypereosinophilic, with cytoplasmic fragmentation and a mild leukocytic infiltrate composed of lymphocytes, granulocytes, and macrophages, indicative of myofiber degeneration and necrosis. HCE stain. Scale bar = 200 µm.

##### Liver, skeletal muscle and gills

Differences in hepatocyte vacuolation was not significantly different between groups (Table 3). Gills had no appreciable abnormalities across the different treatment groups

### 3.3 Weight

Terminal fish weights varied significantly among treatment groups at 50-days post-Dex exposure (Table 3). Control fish averaged 21.5 g (95% CI: 17.9-25.2), while ASE Only fish weighed 19.6 g (95% CI: 14.4-24.7). No significant difference was observed between ASE Only and Control groups. Fish treated with Dex and fasted (DexFast) showed substantially lower weights, averaging 8.72 g (95% CI: 7.01-10.4). DexFast-treated fish had significantly lower weights compared to both Control (diff = -11.7 g, *p* = 0.0002) and ASE Only (diff = - 12.295 g, *p* < 0.0001) treatment groups. Fish in the combined DexFast + ASE treatment group were not weighed as they did not survive to the 50-day sampling point, the timepoint that the fish were weighed. In summary, fasting treatment had a profound effect on fish weight, with the DexFast treatment group resulting in dramatic reductions in body weight (approximately 59% lower body weight than controls). ASE treatment alone had minimal impact on growth with only an 8.8% reduction in body weight compared to controls.

### 3.4 Electron microscopy

#### Field Sample 201G

Transmission electron microscopy (TEM) of enterocytes from the mid-intestine of an ASE-affected, spawning adult spring Chinook Salmon collected in 2019 revealed the presence of putative viral particles (Figure 14). These tissues were previously analyzed for *E. schreckii* (Couch et al., 2022), but the viral particles are newly described from the intestinal tissue of that study. Viruses were observed in one of four fish processed for electron microscopy. The particles measured 90–140 nm in diameter and most exhibited electron-dense cores, an electron-lucent intermediary layer, and a thin, electron-dense outer layer (Figure 14).

**Figure 14.**
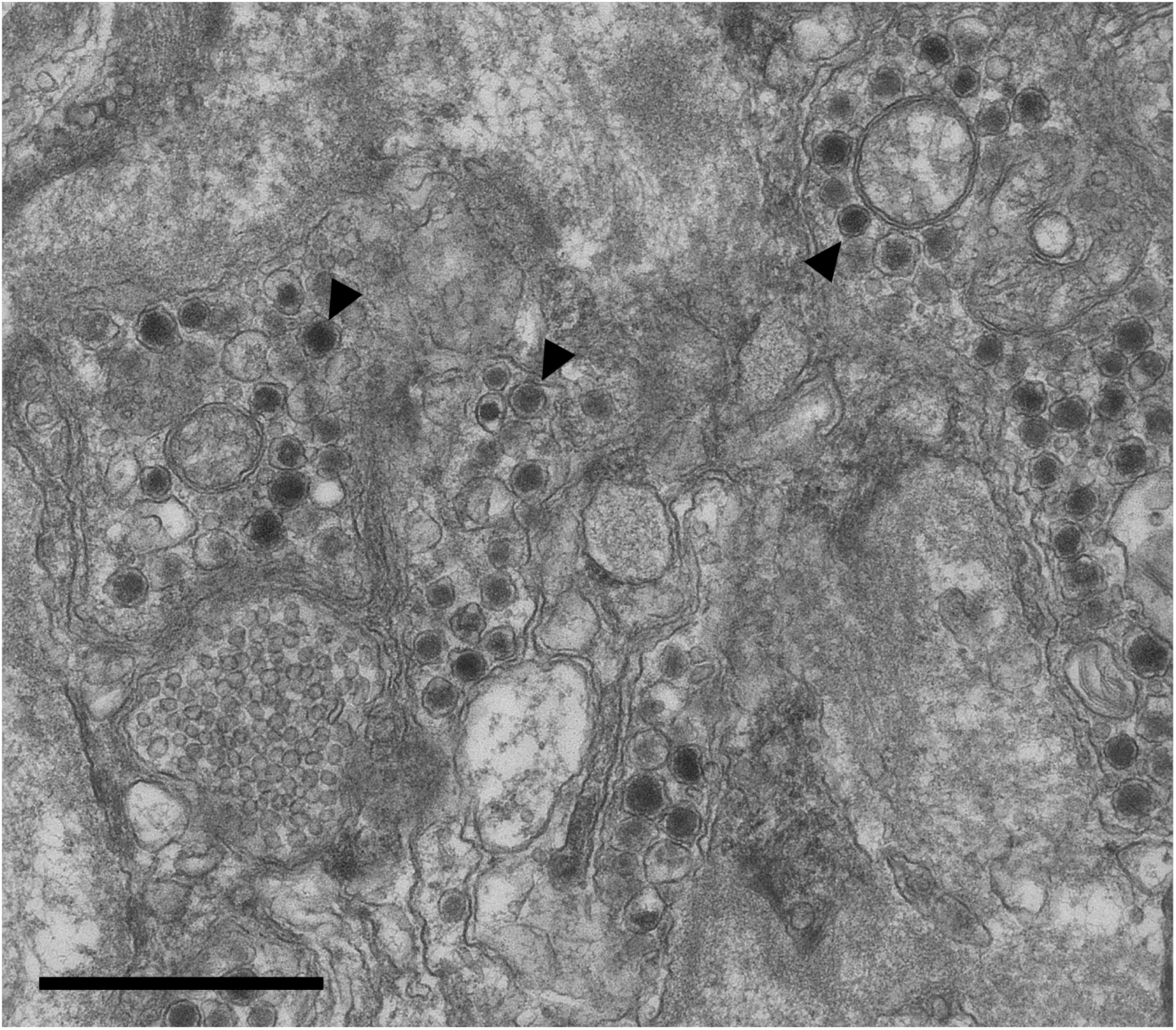
**Transmission electron microscopy (TEM)** of an enterocyte from the mid-intestine of an ASE-affected, spawning adult spring Chinook Salmon collected in 2019. Samples were previously analyzed for *Enterocytozoon schreckii* (Couch et al., 2022), but the putative viral particles (arrowheads) are newly described in this study. These intracytoplasmic particles, measuring 90-140 nm in diameter, exhibit electron-dense cores, an electron-lucent intermediary layer, and a thin, electron-dense outer layer. Scale bar = 1 µm.

#### CLEM: Phase 2

To further characterize the intranuclear inclusions and the presence of *E. schreckii*, we performed correlative light and electron microscopy (CLEM) on enterocytes from the mid-intestine of ASE-inoculum recipient fish. At lower magnification, intranuclear inclusions previously identified by histology appeared as electron-dense structures containing numerous putative viral particles (Figure 15A). Higher magnification revealed that these putative viral particles were pleomorphic, with diameters ranging from 90–115 nm. Notably, the viral particles exhibited a potential lipid bilayer, consisting of two electron-dense layers separated by a thicker electron-lucent layer (Figure 15B). Additionally, aggregates of *E. schreckii* spores were observed within the cytoplasm of the enterocytes (Figure 15C). At higher magnification, the spores exhibited distinct coiled polar filaments, a diagnostic feature of microsporidia (Figure 15D).

**Figure 15.**
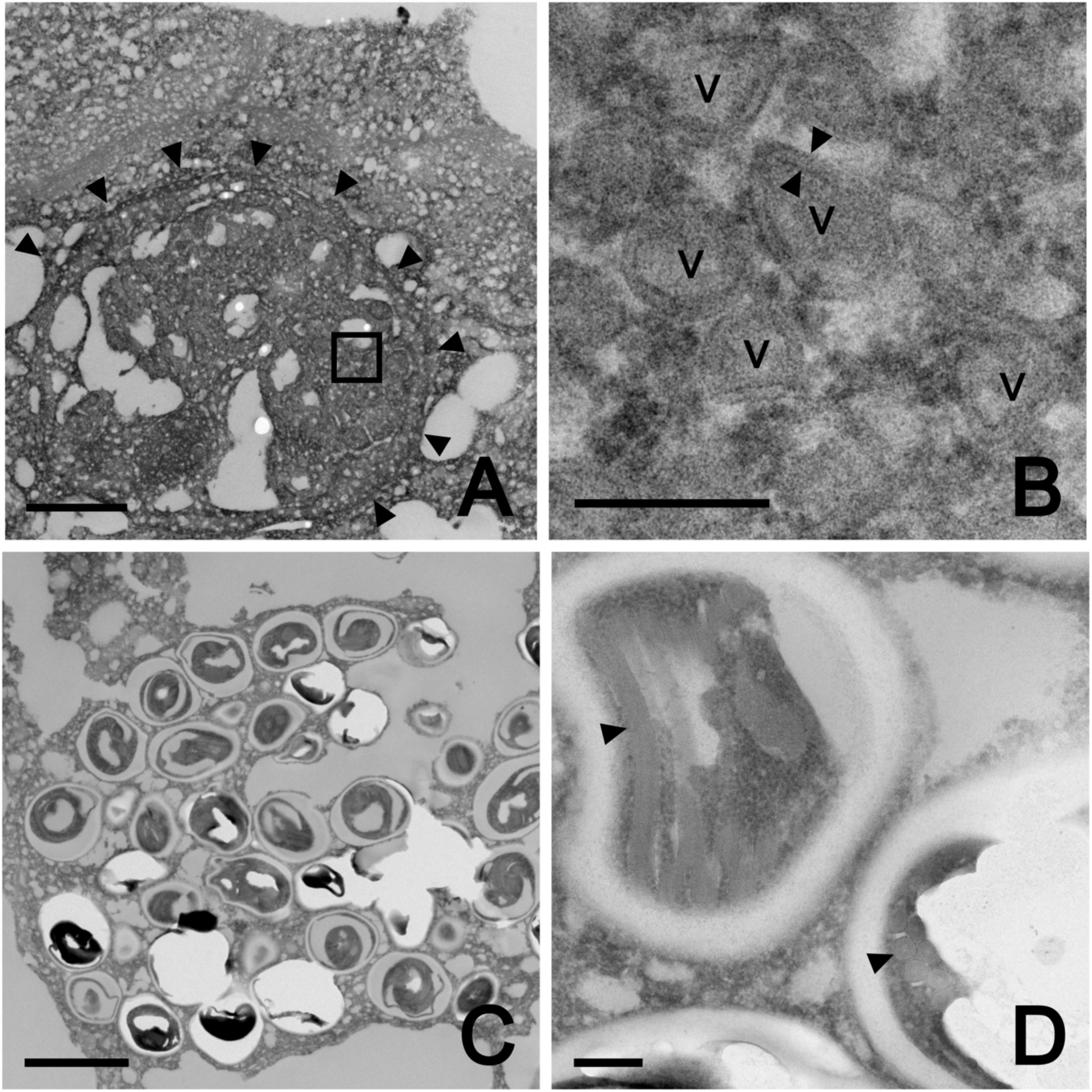
Phase 2 Mid-intestine. Representative Correlative Light Electron Microscopy (CLEM) of enterocytes from the mid-intestine of an ASE-inoculum recipient fish that exhibited viral-like intranuclear inclusions and *Enterocytozoon schreckii* by light microscopy (Figure 8). A) Low magnification of the epithelium reveals an electron-dense inclusion (arrowheads). Scale bar = 2 µm. B) Higher magnification from the boxed area in A of the inclusion highlights the viral particles (V). These particles range from 90-115 nm in diameter. They exhibit a pleomorphic shape and potential lipid bilayer, characterized by two electron-dense layers separated by a thicker electron-lucent layer (arrowheads). Scale bar = 200 nm. C) Aggregate of *E. schreckii* spores within an enterocyte. Scale bar = 2 µm. D) Higher magnification showing spores with coiled polar filaments (arrowheads). Scale bar = 200 nm.

## 4 Discussion

### 4.1 Background

Evidence accumulated over 15 years of research has revealed a correlation between PSM and unique intestinal lesions in Willamette River Chinook Salmon (Couch et al., 2022; Nervino et al., 2024). Here, we define this disease as Adult Salmon Enteritis (ASE) based on the prominent loss of normal epithelium and profound, regionally diffuse inflammation in the intestinal tract (i.e., severe ulcerative enteritis). Thus far, we have only observed this presentation in adult spring Chinook Salmon after they have returned to freshwater to spawn. Nervino et al. (2024) conducted a retrospective histologic review of hundreds of salmon intestines collected over ten years, which included apparently healthy summer fish, PSM fish, successfully spawned fish at hatcheries, and post-spawned fish from the Willamette River and its tributaries. Excluding PSM fish, an expected progression of severity of ASE develops through the summer in certain rivers in Oregon, where many of the artificially spawned fish have severe disease and fish that naturally spawn a few weeks later have similarly severe disease. PSM fish, collected in the summer, have a very similar ASE severity as post-spawned fish, and hence Nervino et al. (2024) showed a strong statistical correlation of severity of ASE (i.e., epithelial loss) with PSM.

We previously considered that ASE may be caused simply by accelerated senescence in this semelparous species due to deleterious environmental conditions or extended time in the river due to returning earlier than expected. However, ASE was completely absent in two populations of successfully spawned Chinook Salmon from Washington State. Due to this lack of ASE in some spring Chinook Salmon populations, along with the severity of the lesions, we have focused on an infectious etiology for the hypothesized cause of ASE and investigated this further by employing laboratory transmission experiments.

### 4.2 Study Results

Our laboratory transmission study demonstrated that juvenile spring Chinook Salmon exposed by oral gavage of intestinal tissues from Chinook Salmon with ASE showed manifestation of intestinal changes consistent with the disease in the recipient fish, particularly in fish that were fasted and immunosuppressed with Dex. Our results here support an impact on mortality of the high dose Dex and fasting treatments, particularly the former, on juvenile Chinook Salmon in both trials, providing a successful model for spawning adult Chinook Salmon in a juvenile fish.

#### 4.2.1 Dexamethasone

Based on mortalities and epizootics by primary and opportunistic infectious agents, we concluded that the fish were immune compromised in both trials, similar to our first report (Couch et al., 2023). Several previous laboratory studies have immunosuppressed salmonid fishes through corticosteroid administration, specifically Dex (Fagerlund & McBride, 1969; Lindenstrøm & Buchmann, 1998; Lovy et al., 2008; McLeay, 1973; Nielsen & Buchmann, 2003; Pickering et al., 1987) and cortisol (McLeay, 1973). For chronic incremental corticosteroid release, IP implants of corticosteroid in an oil vehicle have been used; with either cortisol (Cortés et al., 2013; Kent & Hedrick, 1987; Maule et al., 1987; Niklasson et al., 2014) or Dex (Lindenstrøm & Buchmann, 1998). Dex compromises circulating leukocytes and the innate immune response, circumventing the adaptive immune response, of rainbow trout (*Oncrohynchus mykiss*) (Lovy et al., 2008; Pickering et al., 1987). We also observed morbidity in Dex treated fish associated with branchitis and opportunistic *I. necator* infections as seen in our earlier study at the same facility (Couch et al., 2023), and similar to the study by Lovy et al. (Lovy et al., 2008) with oral Dex administration.

Phase 1 showed that both Dex treatments, low dose and high dose, increased the risk of mortality, where high dose Dex increased the mortality risk by 7-fold compared to control fish. Nevertheless, survival was adequate for two months to allow us to move forward with a similar protocol for Phase 2 experiment. With the latter, fish treated with DexFast also showed high mortality. Dex treatments in salmon have demonstrated increased mortality in similar studies, with increased mortality shown by Lovy et al. (Lovy et al., 2008) caused by chronic exposure via oral administration.

#### 4.2.2 Fasting

Previous studies demonstrated that juvenile salmonids can be fasted without significant impact on mortality for up to 60 days (Mizuno et al., 2002) or even 120 days (Karatas et al., 2021; Pottinger et al., 2003). Our results further support these studies, where the fasting treatments, both low fast and high fast, demonstrated minimal impact on the mortality outcome during the two-month experimental timeframe. Fish that were on the high fast diet (complete fasting) had a 10-15% mortality rate, which was similar to mortality rate reported in masu salmon (*Oncorhynchus masou*) after a 60 day fasting period (Mizuno et al., 2002). We did not weigh fish at the end of this “range finding” experiment with Phase 1, but subjective observations were consistent with the diets, i.e., those fed at 3%/day were larger, those on the 1% maintenance diet were about the same size as the beginning of the experiment, and those receiving no food were smaller.

Fasting has documented effects on intestinal morphology in other salmonids, including reduction of enterocyte height and number, intestinal fold length, and epithelial surface area in juvenile Caspian salmon (*Salmo trutta caspius*) (Emadi Shaibani et al., 2013) and, with mammals, increased individual enterocyte apoptosis in C57BL6 mice (Chappell et al., 2003). Our fasted fish (including fish without Dex treatment in Phase 1) demonstrated similar changes: visually decreased enterocyte size and decreased number of goblet cells compared to control fish based on histopathologic examination (Figure 4), supporting expected fasting effects.

We also examined histopathologic changes in the liver and kidney associated with fasting. Melanin deposition in melano-macrophages within the renal interstitium is an indicator of the degree of starvation/negative energy balance experienced in salmon species (Agius & Roberts, 1981; Mizuno et al., 2002), where melanin increases as the negative energy balance and subsequent catabolism increase. A mild amount of melanin deposition in the kidneys of salmonids is also considered normal, as it was noted by Mizuno et al. (2002) in salmon on a calorically complete diet. Our findings of significantly increased melanin deposition in both low and high fasted fish compared to unfasted fish further supports these findings in Phase 1. Melanin content within the renal interstitial space was increased in the DexFast Only group compared to the ASE Only group in Phase 2, which is most likely due to the fasting diet. Melanin deposition is associated with a lower body condition factor (weight / body length) in salmonids (Schwindt et al., 2006), which can be comparable to weight. To support this hypothesis, the fasting treatment in the DexFast Only group was significantly lower than the Control group – i.e., had 59% lower body weight (Table 3).

Regarding the liver, vacuolar changes in hepatocytes were noted by histopathology in both the Phase 1 and Phase 2 experiments (Figure 7). At severe stages of emaciation, most animals demonstrate atrophy of hepatocellular cytoplasm (Jubb et al., 2016; Pfeifer, 1972; Tongiani, 1971), including jack mackerel larvae (*Trachurus symmetricus*) (Theilacker, 1978) and zebrafish (*Danio rerio*) in late stages of cachexia called “Starvation Syndrome” (https://zebrafish.org/wiki/health/disease_manual/start). We did not observe this severe liver change, indicating that the two months of fasting was not significant enough time to induce prominent emaciation-associated changes in juvenile Chinook Salmon. This is further supported by the consistent presence of adipose tissue, especially visceral adipose within the coelomic cavity of all fish. Hence, these findings support the current fasting regime in our model inducing a catabolic state without eliciting profound starvation-associated changes.

Inflammatory changes observed in the coelom were very likely the result of the lipid-based delivery of the Dex. Chronic inflammation with lipid droplets leading to formation of adhesions is a common side effect in Atlantic salmon (*Salmo salar*) that are immunized by IP oil-adjuvant vaccines for *Aeromonas salmonicida* (Poppe & Breck, 1997). However, this was not noted in other studies using similar delivery of corticosteroids in salmonids (Cortés et al., 2013; Couch et al., 2023; Fagerlund & McBride, 1969; Kent & Hedrick, 1987; Lindenstrøm & Buchmann, 1998; McLeay, 1973; Nielsen & Buchmann, 2003). The absence of significant mortality in control fish, both with and without the sham implant vehicle (Tanks 5 and 6), demonstrated that the handling and injection of the sham implant had minimal impact. This further emphasizes the significant impacts of the Dex treatment on the fish in Tanks 1-4.

#### 4.2.3 Transmission of ASE and pathogens

Exposure to ASE tissue inoculum by oral gavage (Phase 2) was consistently associated with negative effects, including increased risk of mortality, increased enteritis scores, dysplasia of the intestinal epithelium, and presence of putative viral intranuclear inclusions and *E. schreckii* in enterocytes. Interestingly, the DexFast + ASE group lacked significantly higher enteritis scores than all other groups. Dex has been documented to decrease the function of the innate immune response while sparing the adaptive immune response in rainbow trout (*O. mykiss*)(Lovy et al., 2008) and Senegalese sole (*Solea senegalensis*)(Salas-Leiton et al., 2012), leading to increased susceptibility to infections in rainbow trout (Lovy et al., 2008), Atlantic salmon (*Salmo salar*) (Nielsen & Buchmann, 2003), common carp (*Cyprinus carpio*)(Nakayama et al., 2017), Crucian carp (*Carassius carassius*) (Qi et al., 2016), and Senegalese sole (Salas-Leiton et al., 2012). Likewise, sexually mature Chinook Salmon in the field also show impaired innate immunity but still produce antibodies until spawning (Dolan et al., 2016). The functional decrease of the innate immune response caused by Dex treatment accounts for the less severe enteritis score in the DexFast + ASE group compared to the changes in the ASE Only fish and the massive inflammation in the lamina propria seen in field sampled adult salmon with ASE (Nervino et al., 2024). Despite the immune compromised status of ASE-affected adult spring Chinook, they still display severe intestinal inflammation. This is consistent with cortisol causing a reduction in the innate immunity, but less so with adaptive immunity (Dolan et al., 2016). This could also be explained in part by the observations that inflammation is seen before the loss of the epithelium with ASE - e.g., fish collected in the late spring or early summer from the Willamette River had noteworthy inflammation while the epithelium was still intact (Nervino et al., 2024). Consistent with this observation, Dolan et al. (2016) showed that innate immunity dramatically declined as the summer progresses, and (Maule, Schrock, Slater, Fitzpatrick, & Schreck (1996) showed that cortisol levels steeply increase as the summer progresses from June to August. Hence these fish are more immunologically competent early in the summer, when lamina propria inflammation develops.

### 4.3 Further evidence of an infectious agent

The similarity of ASE lesions amongst adult broodstock Chinook Salmon from Oregon rivers (Nervino et al., 2024) and juvenile Chinook exposed to ASE tissue provide strong evidence that these lesions (i.e., ASE) are caused by a transmissible agent. This is consistent with the geographic clustering and epidemic nature of ASE outbreaks observed in sexually mature spring Chinook Salmon in Oregon and Washington, where ASE is profound in some populations and completely absent in others.

The similarities to early lesions in adult broodstock include progressive mixed enteritis and the presence of the microsporidian parasite *E. schreckii*, also seen in Phase 2 fish exposed to ASE. With the additional discovery of putative viral inclusions, consistent with a cytomegalo-like virus, within the nuclei of affected enterocytes, our study provides a viable model to further investigate the underlying causes and mechanisms of ASE or other pathologies that may impact spawning spring Chinook Salmon. *In vivo* transmission studies are valuable for interrogating the possible infectious etiologies of diseases that have clear case definitions, but a specific pathogen is unapparent. For example, Theiler’s disease in horses, historically associated with equine-origin biologics, remained enigmatic until experimental *in vivo* transmission studies identified Equine Parvovirus-Hepatitis (EqPV-H) as a probable cause. In this case, researchers inoculated healthy horses with serum from affected individuals and observed the development of hepatitis, successfully linking EqPV-H to clinical cases of Theiler’s disease and emphasizing the role of viral agents in similar unexplained syndromes (Divers et al., 2022).

The identification of novel diseases often begins with characteristic pathological lesions, but transmission trials are essential to confirm an infectious etiology. Some fish diseases characterized by distinct inflammatory lesions, such as strawberry disease in rainbow trout, were successfully transmitted under laboratory conditions, supporting the role of infectious agents in their etiology (Verner-Jeffreys et al., 2008). Similarly, plasmacytoid leukemia of Chinook Salmon has been studied through transmission trials to confirm its infectious potential (Eaton & Kent, 1992; Kent & Dawe, 1990). In zebrafish, gastrointestinal tumors were transmitted in the laboratory by exposure to water from affected fish and were linked to *Mycoplasma* sp. and *Pseudocapillaria tormentosa* infections (Kent et al., 2021).

### 4.4 Parasites and Viruses

Two intestinal parasites, *C. shasta* and *E. schreckii*, are commonly observed in certain stocks of Oregon spring Chinook Salmon with ASE, but a retrospective evaluation of over 700 intestines of these fish found that neither parasite was statistically correlated with these the disease (Nervino et al., 2024). Moreover, many fish showed ASE without detection of these parasites. *Ceratonova shasta* requires an alternate host (annelid worm) to complete its life cycle (Bartholomew et al., 1997). Consistent with this, all the donor fish contributing to the inoculum in Phase 2 were infected with *C. shasta*, but this myxozoan was not observed in recipient fish.

The microsporidium *Enterocytozoon schreckii* is particularly noteworthy for its direct transmission via the fecal-oral route in Chinook Salmon, and its potential synergistic relationship with viruses in the development of ASE. The microsporidium was detected in 5 of 15 assessed donor broodstock used for the inoculum, and the parasite was subsequently detected in recipient fish in Phase 2, demonstrating that an intermediate host is not required. The microsporidium is of particular interest as natural infections have only been documented in adult salmon populations with ASE (Couch et al., 2022). Moreover, *E. schreckii* shares remarkable similarities with *E. bieneusi*, a clinically significant opportunistic pathogen of immune compromised human patients (i.e., AIDS)(Michiels et al., 1993; Orenstein et al., 1992). *E. bieneusi* infection results in a diarrhea and negative energy balance (Michiels et al., 1993; Orenstein et al., 1992; Weiss & Reinke, 2022). Both microsporidia are genetically similar and infect enterocytes (Couch et al., 2022; Weiss & Reinke, 2022). Infection by *Nucleospora* (syn. *Enterocytozoon*) *salmonis*, a relative of *E. schreckii* and *E. bieneusi* in the family Enterocytozooidae, has been experimentally transmitted in Chinook Salmon through the feeding of infected tissues from the kidney and spleen of infected fish (Baxa-Antonio et al., 1992; Hedrick et al., 1991). Microsporidia of fishes are usually transmitted by the oral ingestion of spores found in the environment, carcasses or in food, resulting in an initial gastrointestinal infection that can then disseminate depending on the species of microsporidia (Kent et al., 2014). Microsporidian invasion of intestinal cells occurs via discharge of the coiled polar filament, a unique microsporidian structure, which injects the sporoplasm directly into host cells where subsequent intracellular development occurs (Sanders et al., 2014; Weiss & Reinke, 2022). Feng et al. (2006) was able to demonstrate the fecal-oral transmission of *E. bieneusi*, using immune-compromised rats and mice, demonstrating how immune suppression facilitates infection and persistence of this relative of *E. schreckii*. This further supports the hypothesis that *E. schreckii* is transmitted between salmonids through fecal-oral routes, and its effects may be magnified in populations with compromised immunity, such as adult spring Chinook Salmon returning to spawn in Oregon.

Whereas most recognized as a parasite of AIDS patients, *E. bieneusi* has been detected in immune competent patients and mammals, typically causing self-limiting diarrhea or remaining asymptomatic (Decraene et al., 2012; Mathis et al., 2005), confirming the potential for asymptomatic carriers and transmission. Likewise, we observed *E. schreckii* in some fish exposed to ASE-affected tissue in Phase 2, despite the absence of immunosuppression with Dex. This raises the possibility that, like *E. bieneusi*, subclinical infections may occur in immunocompetent fish and could be detected using more sensitive testing modalities such as PCR (Kent et al., 2014). Chinook Salmon may acquire these infections as either juveniles in freshwater or later in the marine environment, before becoming anorexic upon their return to freshwater.

#### Viruses

Intranuclear inclusions were observed in the enterocytes in two recipient fish. These appeared similar to cytomegalovirus (Herpesviridae) inclusions in enterocytes of humans with HIV-infection (AIDS) (Griffiths & Reeves, 2021). While cytomegalovirus is not known to infect fish, there are other herpesviruses seen in fish and these may cause intranuclear inclusions (Hanson et al., 2011; Sano, 1995). CLEM revealed putative viral particles within the histologically identified intranuclear inclusions in one of the ASE-exposed study fish (Figure 15A,B), but unlike typical herpesviruses of fishes we did not observe electron dense cores in the viruses. *Tilapia tilapinevirus* (Tilapia Lake Virus) from the family Amnoonviridae, forms inclusions and are somewhat pleomorphic as seen in the present study (Eyngor et al., 2020; He et al., 2023), but without molecular data the observed virus cannot be assigned to any specific group at this time.

It is also plausible that *E. schreckii* and the viral agent act synergistically in ASE pathogenesis or that infection with one organism facilitates infection with the other. Microsporidia are obligate intracellular parasites that exploit immunocompromised hosts and have a broad host range (Weiss & Reinke, 2022), including fishes (Kent et al., 2014). For example, Lovy et al. (2008) demonstrated that immunosuppression with Dex increased the severity of *Loma salmonae* in rainbow trout (*Oncorhynchus mykiss*). Conversely, microsporidia themselves can suppress the immune system and disrupt normal host immune responses (Bernal et al., 2016; Flores et al., 2021; Han et al., 2020; Moretto et al., 2012; Wasson & Peper, 2000), as seen in Chinook Salmon infected with *Nucleospora* (syn. *Enterocytozoon) salmonis* (Wongtavatchai et al., 1995). This immunosuppression could create a favorable environment for secondary infections, including those caused by viruses. However, there is no published evidence that microsporidia can directly transmit a viral pathogens, unlike what is suspected with pinworm (*Enterobius vermicularis*) eggs transmitting the flagellate *Dientamoeba fragilis* (Johnson et al., 2004) or the nematode *Heterakis gallinarum* eggs transmitting the flagellate *Histomonas meleagridis* in turkeys (McDougald, 2005). Viruses have been detected in a variety of parasitic protozoan taxa (e.g., *Leishmania* spp., *Giardia duobedenalis*, *Cryptosporidium* spp.) and may alter the pathogenesis of protozoan infections in their mammalian hosts (Barrow et al., 2020). While microsporidia may indirectly facilitate viral infections through their immunosuppressive effects on the host, it is intriguing—although highly speculative—to consider whether their mechanism of injecting themselves into host cells could also serve as a pathway for viral transmission.

### 4.5 Conclusion

This laboratory transmission study provides evidence that ASE is caused by an infectious agent and is exacerbated in immune compromised and/or fasted fish, mirroring its presentation in adult fish from rivers and hatcheries (Nervino et al., 2024). Electron and light microscopy findings suggest that the observed enteric virus should be considered as a potential pathogen for this unique disease. Additionally, the presence of *E. schreckii* in recipient fish with ASE raises the possibility that this microsporidium plays a role—whether as a primary cause, a secondary opportunist, or even a vector for another pathogen. Given that many microsporidia are small and easily overlooked in HCE-stained intestinal sections, PCR-based diagnostics and specialized histologic stains (i.e., Luna, Giemsa) would help clarify its true prevalence in both laboratory and wild fish populations. However, the altered bacterial microbiome in fish with ASE (Couch et al., 2023) suggests a bacterial component has not been excluded. Ritz et al. (2024) showed that the virome significantly reshapes bacterial communities during stress responses. Conversely, these microbiome changes could also stem from physiologic differences in affected fish due to other stressors, such as environmental factors or non-bacterial pathogens. These interactions extend to co-infections by known pathogens and would be particularly pertinent in spawning adult salmon given their altered immune state and the long list of pathogens that we have observed in adult spring Chinook Salmon (Benda et al., 2015). There are many examples of this phenomenon in fishes (Okon et al., 2023). With salmonids, there are many examples, three are of underlying infections by metacercaria are *Nanophyetus salmincola* associated with more severe bacterial infections (Roon et al., 2015) and mortality by Infectious Hematopoietic Necrosis virus being exacerbated by co-infections with sea lice (*Lepeophtheirus salmonis*) (Long et al., 2019), and metacercaria from different digenean species interacting with each other (Ferguson et al., 2012).

Further studies are needed to clarify and define the infectious etiology of ASE. Laboratory transmission experiments should be repeated, ideally using much larger, desmolted Chinook Salmon to improve survival rates and enhance the robustness of our model. Although unlikely, future experiments should include control inocula from fish unaffected by ASE to rule out the possibility that ASE-like pathology simply results from the introduction of foreign material into the stomach. Additionally, studies using size-filtered inocula to exclude non-viral pathogens should be conducted. Ultimately, a multifaceted approach will be essential to unraveling the complexities of ASE that is associated with PSM of adult spring Chinook Salmon from Oregon. This should include integrating refined experimental models, tissue culture, advanced molecular techniques such as bacterial and viral microbiome analyses, and target PCR testing of fish from the laboratory and field. These studies may lead to viable options for mitigating the catastrophic loses of this iconic species from Oregon.

## Acknowledgements

We thank Drs. Corbin Schuster and Jayde Ferguson for manuscript review and recommendations. We also thank Ms. Olivia Hakanson Department of Fisheries, Wildlife and Conservation and Ms. Ruth Milston-Clements, Department of Microbiology for their assistance with fish acquisitions and husbandry, and Ms. Leslie Cummins, Albert Einstein College of Medicine for processing CLEM samples. The Oregon Cooperative Fish and Wildlife Research Unit is jointly sponsored by the U.S. Geological Survey, the U.S. Fish and Wildlife Service, the Oregon Department of Fish and Wildlife, Oregon State University, and the Wildlife Management Institute. Any use of trade, product, or firm names is for descriptive purposes only and does not imply endorsement by the U.S. Government.

## Data Availability Statement

Provided at https://datadryad.org/stash

## Conflict of Interest Statement

The authors declare that there are no known competing financial interests or personal relationships that could have influenced the work reported in this study.

## Notes

https://doi.org/10.5061/dryad.wwpzgmswq

### Competing Interest Statement

The authors have declared no competing interest.

## References

Agius, C., & Roberts, R. J. (1981). Effects of starvation on the melano-macrophage centres of fish. Journal of Fish Biology, 19(2), 161–169. 10.1111/j.1095-8649.1981.tb05820.x

Barrow, P., Dujardin, J. C., Fasel, N., Greenwood, A. D., Osterrieder, K., Lomonossoff, G., Fiori, P. L., Atterbury, R., Rossi, M., & Lalle, M. (2020). Viruses of protozoan parasites and viral therapy: Is the time now right? Virology Journal, 17(1), 142. 10.1186/s12985-020-01410-1

Bartholomew, J., Whipple, M., Stevens, D., & Fryer, J. (1997). The life cycle of *Ceratomyxa shasta*, a myxosporean parasite of salmonids, requires a freshwater polychaete as an alternate host. The Journal of Parasitology, 83, 859–868. 10.2307/3284281

Baxa-Antonio, D., Groff, J. M., & Hedrick, R. P. (1992). Experimental horizontal transmission of *Enterocytozoon salmonis* to Chinook salmon, *Oncorhynchus tshawytscha*. The Journal of Protozoology, 39(6), 699–702. 10.1111/j.1550-7408.1992.tb04451.x

Benda, S. E., Naughton, G. P., Caudill, C. C., Kent, M. L., & Schreck, C. B. (2015). Cool, Pathogen-Free Refuge Lowers Pathogen-Associated Prespawn Mortality of Willamette River Chinook Salmon. Transactions of the American Fisheries Society, 144(6), 1159–1172. 10.1080/00028487.2015.1073621

Bernal, C. E., Zorro, M. M., Sierra, J., Gilchrist, K., Botero, J. H., Baena, A., & Ramirez-Pineda, J. R. (2016). *Encephalitozoon intestinalis* inhibits dendritic cell differentiation through an IL-6-dependent mechanism. *Frontiers in Cellular and Infection Microbiology*, C. 10.3389/fcimb.2016.00004

Bowerman, T. E., Keefer, M. L., & Caudill, C. C. (2021). Elevated stream temperature, origin, and individual size influence Chinook salmon prespawn mortality across the Columbia River Basin. Fisheries Research, 237, 105874. 10.1016/J.FISHRES.2021.105874

Carey, K. C., Kent, M., Schreck, C. B., Couch, C. E., Whitman, L., & Peterson, J. T. (2024). Influence of stream temperature and human disturbance on prespawn mortality of Chinook Salmon in the Willamette River basin. North American Journal of Fisheries Management, 44(5), 1147–1164. 10.1002/nafm.11035

Chappell, V. L., Thompson, M. D., Jeschke, M. G., Chung, D. H., Thompson, J. C., & Wolf, S. E. (2003). Effects of incremental starvation on gut mucosa. Digestive Diseases and Sciences, 48(4).

Colvin, M. E., Peterson, J. T., Kent, M. L., & Schreck, C. B. (2015). Occupancy modeling for improved accuracy and understanding of pathogen prevalence and dynamics. PLOS ONE, 10(3), e0116605. 10.1371/JOURNAL.PONE.0116605

Cortés, R., Teles, M., Trídico, R., Acerete, L., & Tort, L. (2013). Effects of cortisol administered through slow-release implants on innate immune responses in rainbow trout (*Oncorhynchus mykiss*). International Journal of Genomics, 2013, 1–7. 10.1155/2013/619714

Couch, C. E., Kent, M. L., Weiss, L. M., Takvorian, P. M., Nervino, S., Cummins, L., & Sanders, J. L. (2022). *Enterocytozoon schreckii* n. Sp. Infects the Enterocytes of Adult Chinook Salmon (*Oncorhynchus tshawytscha*) and May Be a Sentinel of Immunosenescence. mSphere. 10.1128/msphere.00908-21

Couch, C. E., Neal, W. T., Herron, C. L., Kent, M. L., Schreck, C. B., & Peterson, J. T. (2023). Gut microbiome composition associates with corticosteroid treatment, morbidity, and senescence in Chinook salmon (*Oncorhynchus tshawytscha*). Scientific Reports 2023 13:1, 13(1), 1–11. 10.1038/s41598-023-29663-0

Decraene, V., Lebbad, M., Botero-Kleiven, S., Gustavsson, A.-M., & Löfdahl, M. (2012). First reported foodborne outbreak associated with microsporidia, Sweden, October 2009. Epidemiology & Infection, 140(3), 519–527. 10.1017/S095026881100077X

Divers, T. J., Tomlinson, J. E., & Tennant, B. C. (2022). The history of Theiler’s disease and the search for its aetiology. The Veterinary Journal, 287, 105878. 10.1016/j.tvjl.2022.105878

Dobbie, I. M. (2019). Bridging the resolution gap: Correlative super-resolution imaging. Nature Reviews Microbiology, 17(6), 337–337. 10.1038/s41579-019-0203-8

Dolan, B. P., Fisher, K. M., Colvin, M. E., Benda, S. E., Peterson, J. T., Kent, M. L., & Schreck, C. B. (2016). Innate and adaptive immune responses in migrating spring-run adult Chinook salmon, *Oncorhynchus tshawytscha*. Fish & Shellfish Immunology, 48, 136–144. 10.1016/J.FSI.2015.11.015

Eaton, W. D., & Kent, M. L. (1992). A retrovirus in Chinook salmon (*Oncorhynchus tshawytscha*) with plasmacytoid leukemia and evidence for the etiology of the disease. Cancer Research, 52(23), 6496–6500.

Emadi Shaibani, M., Mojazi Amiri, B., & Khodabandeh, S. (2013). Starvation and refeeding effects on pyloric caeca structure of Caspian salmon (*Salmo trutta caspius*, Kessler 1877) juvenile. *Tissue and Cell*, *45*(3), 204–210. 10.1016/J.TICE.2013.01.001

Eyngor, M., Zamostiano, R., Kembou Tsofack, J. E., Berkowitz, A., Bercovier, H., Tinman, S., Lev, M., Hurvitz, A., Galeotti, M., Bacharach, E., & Eldar, A. (2020). Identification of a novel RNA virus lethal to tilapia. Journal of Clinical Microbiology, 52(12), 4137–4146. 10.1128/jcm.00827-14

Fagerlund, U. H. M., & McBride, J. R. (1969). Suppression by dexamethasone of interrenal activity in adult sockeye salmon (*Oncorhynchus nerka*). General and Comparative Endocrinology, 12(3), 651–657. 10.1016/0016-6480(69)90186-5

Feng, X., Akiyoshi, D. E., Sheoran, A., Singh, I., Hanawalt, J., Zhang, Q., Widmer, G., & Tzipori, S. (2006). Serial propagation of the microsporidian *Enterocytozoon bieneusi* of human origin in immunocompromised rodents. Infection and Immunity, 74(8), 4424–4429. 10.1128/IAI.00456-06

Ferguson, J. A., Romer, J., Sifneos, J. C., Madsen, L., Schreck, C. B., Glynn, M., & Kent, M. L. (2012). Impacts of multispecies parasitism on juvenile coho salmon (*Oncorhynchus kisutch*) in Oregon. Aquaculture, 3C2–3C3, 184–192. 10.1016/j.aquaculture.2011.07.003

Flores, J., Takvorian, P. M., Weiss, L. M., Cali, A., & Gao, N. (2021). Human microsporidian pathogen *Encephalitozoon intestinalis* impinges on enterocyte membrane trafficking and signaling. Journal of Cell Science, 134(5), jcs253757. 10.1242/jcs.253757

Grambsch, P. M., & Therneau, T. M. (1994). Proportional hazards tests and diagnostics based on weighted residuals. Biometrika, 81(3), 515–526.

Griffiths, P., & Reeves, M. (2021). Pathogenesis of human cytomegalovirus in the immunocompromised host. Nature Reviews Microbiology, 19(12), 759–773. 10.1038/s41579-021-00582-z

Han, Y., Gao, H., Xu, J., Luo, J., Han, B., Bao, J., Pan, G., Li, T., & Zhou, Z. (2020). Innate and adaptive immune responses against microsporidia infection in mammals. Frontiers in Microbiology, 11, 1468. 10.3389/fmicb.2020.01468

Hanson, L., Dishon, A., & Kotler, M. (2011). Herpesviruses that infect fish. Viruses, 3(11), 2160–2191. 10.3390/v3112160

He, T., Zhang, Y.-Z., Gao, L.-H., Miao, B., Zheng, J.-S., Pu, D.-C., Zhang, Q.-Q., Zeng, W.-W., Wang, D.-S., Su, S.-Q., & Zhu, S. (2023). Identification and pathogenetic study of tilapia lake virus (TiLV) isolated from naturally diseased tilapia. Aquaculture, 5C5, 739166. 10.1016/j.aquaculture.2022.739166

Hedrick, R. P., Groff, J. M., & Baxa, D. V. (1991). Experimental infections with *Enterocytozoon salmonis* Chilmonczyk, Cox, Hedrick (Microsporea): An intranuclear microsporidium from Chinook salmon *Oncorhynchus tshawytscha*. Diseases of Aquatic Organisms, 10, 103–108. 10.3354/dao010103

Johnson, E. H., Windsor, J. J., & Clark, C. G. (2004). Emerging from obscurity: Biological, clinical, and diagnostic aspects of *Dientamoeba fragilis*. Clinical Microbiology Reviews, 17(3), 553–570. 10.1128/cmr.17.3.553-570.2004

Jubb, Kennedy, & Palmer. (2016). Pathology of domestic animals: Volume 2. In M. Grant Maxie (Ed.), Jubb, Kennedy & Palmer’s Pathology of Domestic Animals (Vol. 2). Elsevier.

Karatas, T., Onalan, S., & Yildirim, S. (2021). Effects of prolonged fasting on levels of metabolites, oxidative stress, immune-related gene expression, histopathology, and DNA damage in the liver and muscle tissues of rainbow trout (*Oncorhynchus mykiss*). Fish Physiology and Biochemistry, 47(4), 1119–1132. 10.1007/s10695-021-00949-2

Keefer, M. L., Taylor, G. A., Garletts, D. F., Gauthier, G. A., Pierce, T. M., & Caudill, C. C. (2010). Prespawn mortality in adult spring Chinook salmon outplanted above barrier dams. Ecology of Freshwater Fish, 19(3), 361–372. 10.1111/J.1600-0633.2010.00418.X

Kent, M. L., Benda, S., St-Hilaire, S., & Schreck, C. B. (2013). Sensitivity and specificity of histology for diagnoses of four common pathogens and detection of nontarget pathogens in adult Chinook salmon (*Oncorhynchus tshawytscha*) in fresh water. Journal of Veterinary Diagnostic Investigation, 25(3), 341–351. 10.1177/1040638713482124

Kent, M. L., & Dawe, S. C. (1990). Experimental transmission of a plasmacytoid leukemia of Chinook salmon, *Oncorhynchus tshawytscha*. Cancer Research, 50(17 Suppl), 5679s–5681s.

Kent, M. L., & Hedrick, R. P. (1987). Effects of cortisol implants on the PKX myxosporean causing proliferative kidney disease in rainbow trout, *Salmo gairdneri*. The Journal of Parasitology, 73(3), 455–461. 10.2307/3282121

Kent, M. L., Shaw, R. W., & Sanders, J. L. (2014). Microsporidia in Fish. In Microsporidia (pp. 493–520). John Wiley & Sons, Ltd. 10.1002/9781118395264.ch20

Kent, M. L., Wall, E. S., Sichel, S., Watral, V., Stagaman, K., Sharpton, T. J., & Guillemin, K. (2021). *Pseudocapillaria tomentosa*, *Mycoplasma* spp., and intestinal lesions in experimentally infected zebrafish *Danio rerio*. Zebrafish, 18(3), 207–220. 10.1089/zeb.2020.1955

Leary, S., Underwood, W., Anthony, R., Cartner, S., Grandin, T., Greenacre, C., Gwaltney-Brant, S., McCrackin, M. A., Meyer, R., Miller, D., Shearer, J., Turner, T., Yanong, R., Johnson, C. L., & Patterson-Kane, E. (2020). AVMA Guidelines for the Euthanasia of Animals: 2020 Edition (pp. 82–91).

Lindenstrøm & Buchmann. (1998). Dexamethasone treatment increases susceptibility of rainbow trout, *Oncorhynchus mykiss* (Walbaum), to infections with *Gyrodactylus derjavini* Mikailov. Journal of Fish Diseases, 21(1), 29–38. 10.1046/j.1365-2761.1998.00070.x

Long, A., Garver, K. A., & Jones, S. R. M. (2019). Synergistic osmoregulatory dysfunction during salmon lice (*Lepeophtheirus salmonis*) and infectious hematopoietic necrosis virus co-infection in sockeye salmon (*Oncorhynchus nerka*) smolts. Journal of Fish Diseases, 42(6), 869–882. 10.1111/jfd.12989

Lovy, J., Speare, D. J., Stryhn, H., & Wright, G. M. (2008). Effects of dexamethasone on host innate and adaptive immune responses and parasite development in rainbow trout *Oncorhynchus mykiss* infected with *Loma salmonae*. Fish & Shellfish Immunology, 24(5), 649–658. 10.1016/j.fsi.2008.02.007

Mathis, A., Weber, R., & Deplazes, P. (2005). Zoonotic potential of the microsporidia. Clinical Microbiology Reviews, 18(3), 423–445. 10.1128/cmr.18.3.423-445.2005

Maule, A. G., Schreck, C. B., & Kaattari, S. L. (1987). Changes in the immune system of Coho salmon (*Oncorhynchus kisutch*) during the parr-to-smolt transformation and after implantation of cortisol. Canadian Journal of Fisheries and Aquatic Sciences, 44(1), 161–166. 10.1139/f87-021

Maule, A. G., Schrock, R., Slater, C., Fitzpatrick, M. S., & Schreck, C. B. (1996). Immune and endocrine responses of adult Chinook salmon during freshwater immigration and sexual maturation. Fish & Shellfish Immunology, 6(3), 221–233. 10.1006/fsim.1996.0022

McDougald, L. R. (2005). Blackhead disease (histomoniasis) in poultry: A critical review. Avian Diseases, 49(4), 462–476. 10.1637/7420-081005R.1

McLeay, D. J. (1973). Effects of cortisol and dexamethasone on the pituitary-interrenal axis and abundance of white blood cell types in juvenile Coho salmon, *Oncorhynchus kisutch*. General and Comparative Endocrinology, 21(3), 441–450. 10.1016/0016-6480(73)90103-2

Michiels, J. F., Hofman, P., Saint Paul, M. C., Loubière, R., Bernard, E., & LeFichoux, Y. (1993). Pathological features of intestinal microsporidiosis in HIV positive patients. Pathology - Research and Practice, 189(4), 377–383. 10.1016/S0344-0338(11)80322-5

Mizuno, S., Misaka, N., Miyakoshi, Y., Takeuchi, K., & Kasahara, N. (2002). Effects of starvation on melano-macrophages in the kidney of masu salmon (*Oncorhynchus masou*). Aquaculture, 209(1–4), 247–255. 10.1016/S0044-8486(01)00716-5

Moretto, M. M., Khan, I. A., & Weiss, L. M. (2012). Gastrointestinal cell mediated immunity and the microsporidia. PLoS Pathogens, 8(7), e1002775. 10.1371/journal.ppat.1002775

Nakayama, K., Yamashita, R., & Kitamura, S.-I. (2017). Use of common carp (*Cyprinus carpio*) and *Aeromonas salmonicida* for detection of immunomodulatory effects of chemicals on fish. Marine Pollution Bulletin, 124(2), 710–713. 10.1016/j.marpolbul.2016.12.060

Naughton, G. P., Keefer, M. L., Clabough, T. S., Knoff, M. J., Blubaugh, T. J., Morasch, M. R., Sharpe, C. S., & Caudill, C. C. (2023). Prespawn mortality of spring Chinook salmon in three Willamette river populations. North American Journal of Fisheries Management, 43(3), 715–729. 10.1002/nafm.10887

Nervino, S., Polley, T., Peterson, J. T., Schreck, C. B., Kent, M. L., & Alexander, J. D. (2024). Intestinal lesions and parasites associated with senescence and prespawn mortality in Chinook Salmon (*Oncorhynchus tshawytscha*). Journal of Fish Diseases, 47(2), e13876. 10.1111/jfd.13876

Nielsen, C. V., & Buchmann, K. (2003). Increased susceptibility of Atlantic salmon *Salmo salar* to infections with *Gyrodactylus derjavini* induced by dexamethasone bath treatment. Journal of Helminthology, 77(1), 65–68. 10.1079/JOH2002159

Niklasson, L., Sundh, H., Olsen, R.-E., Jutfelt, F., Skjødt, K., Nilsen, T. O., & Sundell, K. S. (2014). Effects of cortisol on the intestinal mucosal immune response during cohabitant challenge with IPNV in Atlantic salmon (*Salmo salar*). PLoS ONE, 9(5), e94288. 10.1371/journal.pone.0094288

Okon, E. M., Okocha, R. C., Taiwo, A. B., Michael, F. B., & Bolanle, A. M. (2023). Dynamics of co-infection in fish: A review of pathogen-host interaction and clinical outcome. Fish and Shellfish Immunology Reports, 4, 100096. 10.1016/j.fsirep.2023.100096

Orenstein, J. M., Tenner, M., & Kotler, D. P. (1992). Localization of infection by the microsporidian *Enterocytozoon bieneusi* in the gastrointestinal tract of AIDS patients with diarrhea. AIDS, 6(2), 195.

Peterson, T., Spitsbergen, J., Feist, S., & Kent, M. (2011). Luna stain, an improved selective stain for detection of microsporidian spores in histologic sections. *Diseases of Aquatic Organisms*, 95(2), 175–180. 10.3354/dao02346

Pfeifer, U. (1972). Cellular autophagy and cell atrophy in the rat liver during long-term starvation: A quantitative morphological study with regard to diurnal variations. Virchows Archiv B Cell Pathology, 12(1), 195–211. 10.1007/BF02893998

Pickering, A. D., Pottinger, T. G., & Sumpter, J. P. (1987). On the use of dexamethasone to block the pituitary-interrenal axis in the brown trout, *Salmo trutta* L. *General and Comparative Endocrinology*, 65(3), 346–353. 10.1016/0016-6480(87)90119-5

Poppe, T., & Breck, O. (1997). Pathology of Atlantic salmon *Salmo salar* intraperitoneally immunized with oil-adjuvanted vaccine. A case report. Diseases of Aquatic Organisms, 29, 219–226. 10.3354/dao029219

Posit team. (2024). RStudio: Integrated development environment for R (Version Kousa Dogwood, 2024.12.0.467) [Computer software]. Posit Software, PBC. http://www.posit.co/

Pottinger, T. G., Rand-Weaver, M., & Sumpter, J. P. (2003). Overwinter fasting and re-feeding in rainbow trout: Plasma growth hormone and cortisol levels in relation to energy mobilisation. Comparative Biochemistry and Physiology Part B: Biochemistry and Molecular Biology, 136(3), 403–417. 10.1016/S1096-4959(03)00212-4

Qi, X.-Z., Li, D.-L., Tu, X., Song, C.-G., Ling, F., & Wang, G.-X. (2016). Preliminary study on the relationship between dexamethasone and pathogen susceptibility on crucian carp (*Carassius auratus*). Fish & Shellfish Immunology, 59, 18–24. 10.1016/j.fsi.2016.10.017

R Core Team. (2024). R: A language and environment for statistical computing (Version R version 4.4.1 (2024-06-14)) [Computer software]. R Foundation for Statistical Computing. https://www.R-project.org/

Ritz, N. L., Draper, L. A., Bastiaanssen, T. F. S., Turkington, C. J. R., Peterson, V. L., van de Wouw, M., Vlckova, K., Fülling, C., Guzzetta, K. E., Burokas, A., Harris, H., Dalmasso, M., Crispie, F., Cotter, P. D., Shkoporov, A. N., Moloney, G. M., Dinan, T. G., Hill, C., & Cryan, J. F. (2024). The gut virome is associated with stress-induced changes in behaviour and immune responses in mice. Nature Microbiology, 9(2), 359–376. 10.1038/s41564-023-01564-y

Roon, S. R., Alexander, J. D., Jacobson, K. C., & Bartholomew, J. L. (2015). Effect of *Nanophyetus salmincola* and bacterial co-infection on mortality of juvenile Chinook salmon. Journal of Aquatic Animal Health, 27(4), 209–216. 10.1080/08997659.2015.1094150

Roumasset, A. (2012). Pre-spawn mortality of Upper Willamette River spring Chinook salmon: Associations with stream temperature, watershed attributes, and environmental conditions on the spawning grounds [Thesis: Master of Science in Water Resources, University of Idaho]. 10.13140/RG.2.2.36796.85129

Salas-Leiton, E., Coste, O., Asensio, E., Infante, C., Cañavate, J. P., & Manchado, M. (2012). Dexamethasone modulates expression of genes involved in the innate immune system, growth and stress and increases susceptibility to bacterial disease in Senegalese sole (*Solea senegalensis* Kaup, 1858). Fish & Shellfish Immunology, 32(5), 769–778. 10.1016/j.fsi.2012.01.030

Sanders, J. L., Peterson, T. S., & Kent, M. L. (2014). Early development and tissue distribution of *Pseudoloma neurophilia* in the zebrafish, *Danio rerio*. Journal of Eukaryotic Microbiology, 61(3), 238–246. 10.1111/jeu.12101

Sano, T. (1995). Viruses and viral diseases of salmonids. Aquaculture, 132(1), 43–52. 10.1016/0044-8486(94)00372-U

Schreck, C. (2000). Accumulation and long-term effects of stress in fish. CABI, 147–158. 10.1079/9780851993591.0147

Schwindt, A., Truelove, N., Schreck, C., Fournie, J., Landers, D., & Kent, M. (2006). Quantitative evaluation of macrophage aggregates in brook trout *Salvelinus fontinalis* and rainbow trout *Oncorhynchus mykiss*. Diseases of Aquatic Organisms, 68, 101–113. 10.3354/dao068101

Sjoberg, D., Baillie, M., Fruechtenicht, C., Haesendonckx, S., & Treis, T. (2024). ggsurvfit: Flexible time-to-event figures (Version 1.1.0) [Computer software]. https://CRAN.R-project.org/package=ggsurvfit

Theilacker, G. H. (1978). Effect of starvation on the histological and morphological characteristics of jack mackerel, Trachurus symmetricus, larvae. Fishery Bulletin, 76(2), 403–414.

Therneau, T. (2024). A package for survival analysis in R (Version R package version 3.7-0) [Computer software]. https://CRAN.R-project.org/package=survival

Therneau, T. M., & Grambsch, P. M. (2000). Modeling survival data: Extending the cox model [Computer software].

Tongiani, R. (1971). Hepatocyte classes during liver atrophy due to starvation in the golden hamster. Zeitschrift for Zellforschung Und Mikroskopische Anatomie, 122(4), 467–478. 10.1007/BF00936081

Verner-Jeffreys, D. W., Pond, M. J., Peeler, E. J., Rimmer, G. S. E., Oidtmann, B., Way, K., Mewett, J., Jeffrey, K., Bateman, K., Reese, R. A., & Feist, S. W. (2008). Emergence of cold water strawberry disease of rainbow trout *Oncorynchus mykiss* in England and Wales: Outbreak investigations and transmission studies. Diseases of Aquatic Organisms, 79(3), 207–218. 10.3354/dao01916

Wasson, K., & Peper, R. L. (2000). Mammalian microsporidiosis. Veterinary Pathology, 37(2), 113–128. 10.1354/vp.37-2-113

Weiss, L. M., & Reinke, A. W. (Eds.). (2022). Microsporidia: Current advances in biology (Vol. 114). Springer International Publishing. 10.1007/978-3-030-93306-7

Wickham, H. (2016). ggplot2: Elegant graphics for data analysis [Computer software]. https://ggplot2.tidyverse.org

Wongtavatchai, J., Conrad, P. A., & Hedrick, R. P. (1995). Effect of the microsporidian *Enterocytozoon salmonis* on the immune response of Chinook salmon. Veterinary Immunology and Immunopathology, 48(3), 367–374. 10.1016/0165-2427(95)05435-9

